# Soluble mannose receptor induces pro-inflammatory macrophage activation and metaflammation

**DOI:** 10.1101/2020.09.29.315598

**Authors:** Maria Embgenbroich, Hendrik J.P. van der Zande, Leonie Hussaarts, Jonas Schulte-Schrepping, Leonard R. Pelgrom, Noemí García-Tardón, Laura Schlautmann, Isabel Stoetzel, Kristian Händler, Joost M. Lambooij, Anna Zawistowska-Deniziak, Lisa Hoving, Karin de Ruiter, Marjolein Wijngaarden, Hanno Pijl, Ko Willems van Dijk, Bart Everts, Vanessa van Harmelen, Maria Yazdanbakhsh, Joachim L. Schultze, Bruno Guigas, Sven Burgdorf

## Abstract

Pro-inflammatory activation of macrophages in metabolic tissues is critically important in induction of obesity-induced metaflammation. Here, we demonstrate that the soluble mannose receptor (sMR) plays a direct, functional role in both macrophage activation and metaflammation. We show that sMR binds CD45 on macrophages and inhibits its phosphatase activity, leading to a Src/Akt/NF-κB-mediated cellular reprogramming towards an inflammatory phenotype both *in vitro* and *in vivo.* Remarkably, increased serum sMR levels were observed in obese mice and humans and directly correlated with body weight. Additionally, MR deficiency lowers pro-inflammatory macrophages in metabolic tissues and protects against hepatic steatosis and whole-body metabolic dysfunctions in high-fat diet-induced obese mice. Conversely, administration of sMR in lean mice increases serum pro-inflammatory cytokines, activates tissue macrophages and promotes insulin resistance. Altogether, our results reveal sMR as novel regulator of pro-inflammatory macrophage activation which could constitute a new therapeutic target for metaflammation and other hyperinflammatory diseases.

## Introduction

Metaflammation defines a chronic inflammatory state in response to prolonged excessive nutrient intake and is characterized by low-grade inflammation of metabolic tissues (Brestoff and Artis, 2015). Macrophage reprogramming towards an inflammatory phenotype plays a critical role in obesity-induced metaflammation (Lackey and Olefsky, 2016; Hotamisligil, 2017). In lean individuals, macrophages in metabolic tissues maintain tissue homeostasis and insulin sensitivity, potentially through secreting anti-inflammatory cytokines, e.g. TGFβ and IL-10 (Brestoff and Artis, 2015). In metaflammation, however, macrophages in adipose tissue and liver are activated through pro-inflammatory factors in their microenvironment, such as high levels of saturated free fatty acids and IFN-γ. Consequently, these macrophages produce high amounts of tumor necrosis factor (TNF), which directly inhibits canonical insulin signaling (Hotamisligil et al., 1994), leading to ectopic fat deposition in the liver and in skeletal muscles (Shulman, 2014). Additionally, activation of Kupffer cells (KCs), the liver-resident macrophages, promotes recruitment and activation of inflammatory monocytes, which contribute to hepatic insulin resistance and steatosis (Lanthier et al., 2010; Neuschwander-Tetri, 2010; Morinaga et al., 2015).

The MR (also termed CD206) is a type I transmembrane protein belonging to the C-type lectin family which is mainly expressed by subpopulations of macrophages dendritic cells and epithelial cells (Takahashi et al., 1998; Martinez-Pomares, 2012). The MR consists of a cysteine-rich region, a fibronectin type II domain, eight C-type lectin-like domains (CTLDs), a transmembrane region and a short cytosolic tail. Due to its high affinity for glycosylated antigens, the MR plays an important role in antigen uptake and presentation (Burgdorf et al., 2006; Burgdorf et al., 2007). In addition to its functions as a transmembrane protein, the extracellular part of the MR can be shed by metalloproteases and released into the extracellular space (Martínez-Pomares et al., 1998; Jordens et al., 1999). Hence, soluble MR (sMR) can be detected in murine and human serum, and its level was found to be increased in patients with a variety of inflammatory diseases (Andersen et al., 2014; Rødgaard-Hansen et al., 2015; Ding et al., 2017; Suzuki et al., 2018; Loonen et al., 2019; Saha et al., 2019), correlating with severity of disease and even mortality. However, a physiological role of the sMR has not been studied yet, and it remains unclear whether the sMR can actively trigger inflammation.

Here, we report that sMR enhances macrophage pro-inflammatory activation, both in vitro and in vivo, and promotes metaflammation. We demonstrate that the sMR directly interacts with CD45 on the surface of macrophages and inhibits its phosphatase activity, leading to Src/Akt/NF-κB-mediated cellular reprogramming towards an inflammatory phenotype. Additionally, we found enhanced sMR serum levels in obese mice and humans, and show that sMR-induced activation of macrophages triggers metaflammation in vivo.

## Results

### Soluble MR enhances pro-inflammatory cytokine secretion by macrophages

To investigate whether the MR is involved in the pro-inflammatory activation of macrophages, we first stimulated bone marrow-derived macrophages from wild-type or MR-deficient mice with LPS. We found increased secretion of the pro-inflammatory cytokines TNF and IL-6 in MR-expressing wild-type macrophages (Figure 1A). Because the MR itself lacks intracellular signaling motifs and hence, no MR-mediated signaling has been described so far, we hypothesized that the sMR, resulting from the shedding of the MR extracellular region (Figure S1A), might play a role in macrophage activation through direct interaction with macrophage surface proteins. To investigate this hypothesis, we generated a fusion protein consisting of the Fc region of human IgG1 and the extracellular region of the MR (encompassing the cysteine-rich region, the fibronectin region and CTLD1-2) (FcMR) (Schuette et al., 2016). We showed that treatment of MR-deficient macrophages with FcMR also enhanced secretion of TNF and IL-6 after LPS stimulation compared to isotype control-treated cells (Figure 1B). We observed similar results when treating MR-deficient macrophages with commercially available recombinant MR protein, consisting of the complete extracellular region of the protein (Figure 1C), suggesting that binding of sMR to the macrophage surface might indeed be responsible for the observed effects. To definitively prove that the sMR causes the observed increase in cytokine production, we purified sMR from the supernatant of MR-expressing macrophages (Figure S1B) and showed that its administration to MR-deficient macrophages increased the secretion of TNF after LPS stimulation (Figure S1C). Similar results were obtained from FcMR-treated primary macrophages isolated from murine liver, spleen or peritoneal cavity (Figure 1D), and from human monocyte-derived macrophages after addition of recombinant human MR (Figure 1E) or siRNA-mediated down-regulation of the MR (Figure 1F), demonstrating that the sMR enhances pro-inflammatory activation of both murine and human macrophages.

**Figure 1:**
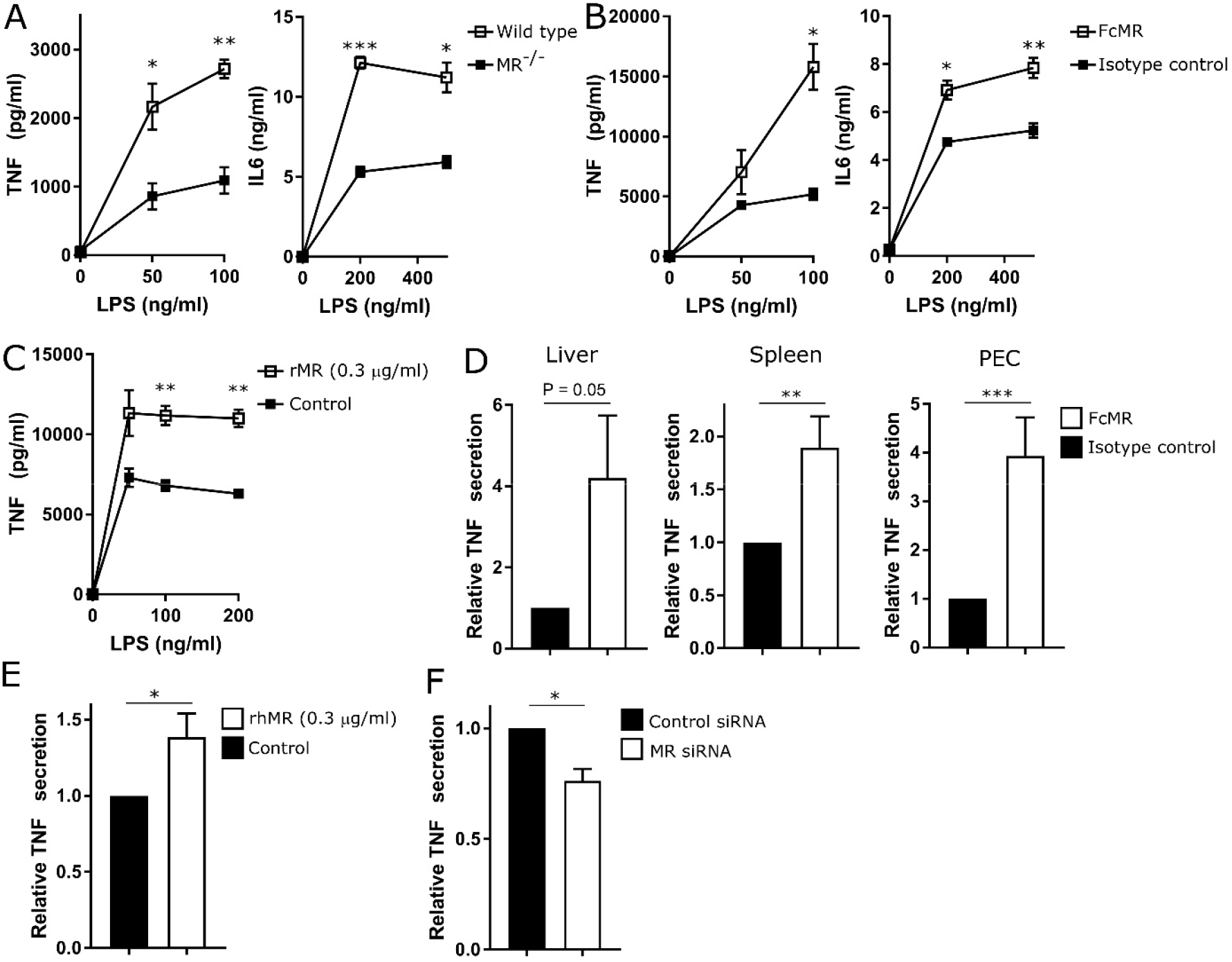
sMR induces pro-inflammatory cytokine secretion by macrophages. A) Secretion of TNF and IL-6 by LPS-treated wild-type or MR-deficient macrophages. B) TNF and IL-6 secretion of LPS-treated MR-deficient macrophages after incubation with FcMR. C) Secretion of TNF by LPS-stimulated MR-deficient macrophages after addition of recombinant MR (rMR). D) Primary murine macrophages were isolated from liver, spleen or peritoneal cavity (PEC) by magnetic separation of F4/80^+^ cells. Secretion of TNF after LPS treatment and stimulation with FcMR were determined by ELISA. E) Secretion of TNF by LPS-treated human monocyte-derived macrophages after stimulation with recombinant human MR (rhMR). F) Secretion of TNF by LPS-stimulated human monocyte-derived macrophages after siRNA-mediated down-regulation of the MR. All graphs are depicted as mean ± SEM; for all experiments, n ≥ 3. See also Figure S1.

### sMR induces a pro-inflammatory phenotype in macrophages

To further dissect the effect of the sMR on macrophages, we treated MR-deficient macrophages with FcMR for 4 h, 12 h or 24 h and performed RNA-seq analysis (Figure 2A). Principle component analysis (PCA) revealed clear transcriptomic distinction of the samples in all analyzed conditions (Figure 2B). A heatmap of the 1366 differentially expressed (DE) genes between FcMR treatment and control presented the substantial changes in gene expression due to the FcMR treatment over time with overlapping and unique gene sets (Figures 2C, S2A). Gene ontology enrichment analysis (GOEA) based on these shared and specific DE gene sets upregulated upon FcMR treatment clearly confirmed inflammatory activation of macrophages (Figure S2B). The most significantly upregulated genes in response to FcMR treatment further emphasized the strong and dynamic pro-inflammatory activation of macrophages, with well-known immunological key mediators such as TNF, IL-6, IL-1β, and IL-12, exhibiting very specific expression patterns (Figure 2D). To classify the response elicited by sMR within the broad spectrum of macrophage activation phenotypes, we performed an enrichment analysis using macrophage activation signatures derived from our previous study comprising macrophages treated with 28 different immunological stimuli (Xue et al., 2014) and the gene sets of FcMR-mediated up-regulated genes per time point. This analysis revealed a striking similarity of FcMR-induced expression patterns to macrophage signatures associated with a chronic inflammatory phenotype, as induced by TNF, PGE2, and P3C (TPP) in our previous stimulation study (Figures 2E, S2C), further substantiating that the sMR reprograms macrophages towards a pro-inflammatory phenotype.

**Figure 2:**
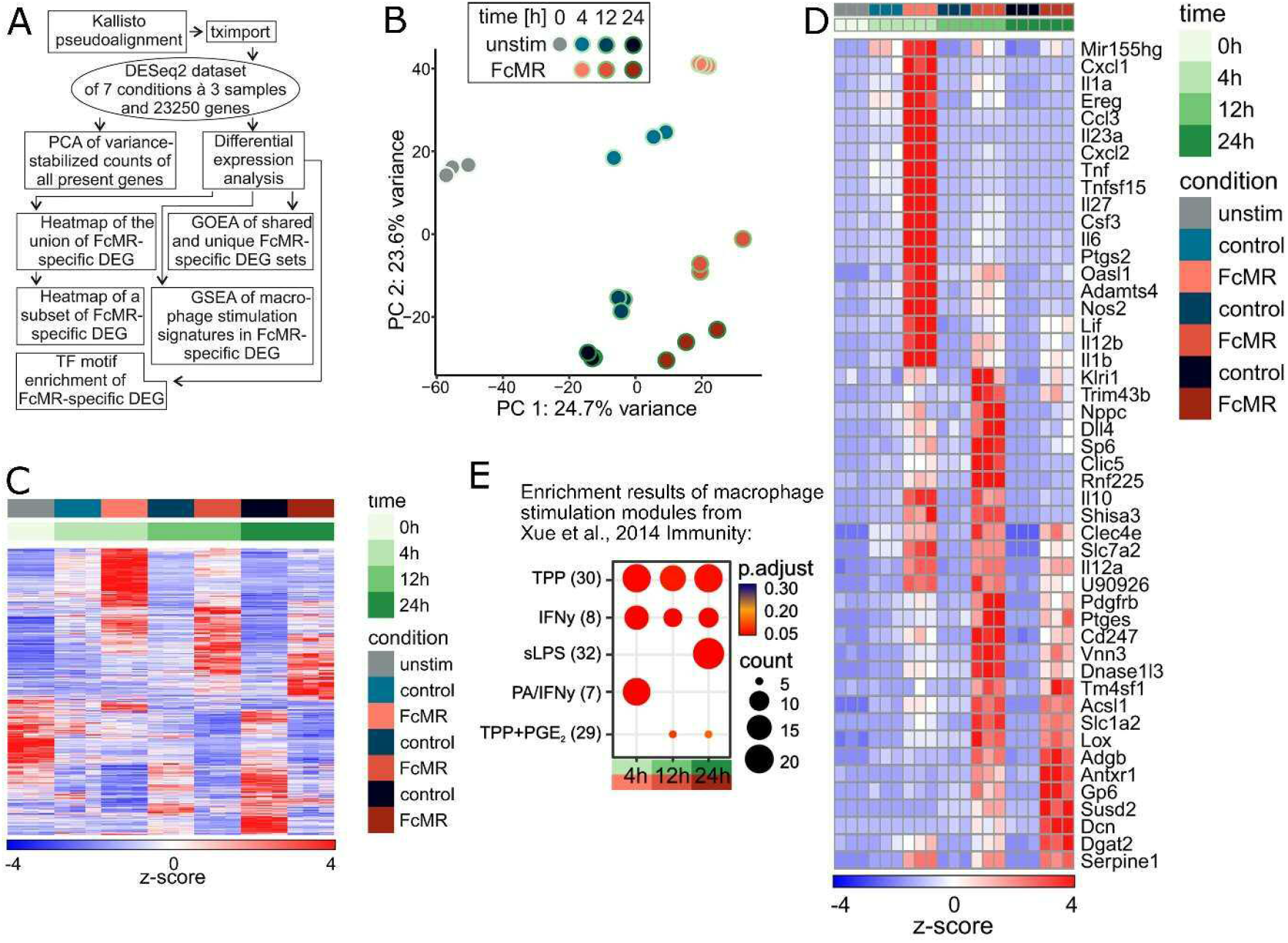
RNAseq analysis of MR-treated macrophages. A) Schematic overview of the bioinformatics RNA-seq analysis strategy. B) Principle component analysis based on variance-stabilized counts of all 23250 present genes. C) Heatmap of hierarchically clustered, normalized and z-scaled expression values of the union of 1366 DE genes between FcMR-treated and control samples. D) Normalized and z-scaled expression values of the union of the top 25 DE genes of each time point significantly upregulated in at least two consecutive time points ranked according to their FcMR vs control samples visualized in a heatmap. E) Dot plot of gene set enrichment analysis results of 49 pre-defined stimulus-specific macrophage expression signatures comprising 28 different stimuli on the FcMR-specific DE genes for each time point. TPP: TNF, PGE2, and Pam3Cys; PA: palmitic acid. See also Figure S2.

### sMR activates macrophages by binding and inhibiting CD45

Next, we investigated the molecular mechanisms regulating sMR-induced macrophage reprogramming and searched for binding partners of the MR on the macrophage surface. To this end, we isolated cell lysates from macrophages that previously underwent surface biotinylation, and performed immunoprecipitation using FcMR. Western Blot analysis using streptavidin allowed us to monitor cell surface proteins interacting with sMR, including a clear band at the molecular weight of the phosphatase CD45 (between 180 and 220 kDa, depending on the splice variant) (Figure S3A), a known binding partner of the MR (Martínez-Pomares et al., 1999). Indeed, co-immunoprecipitation experiments revealed a physical interaction between the MR and CD45 on macrophages (Figures 3A, 3B).

**Figure 3:**
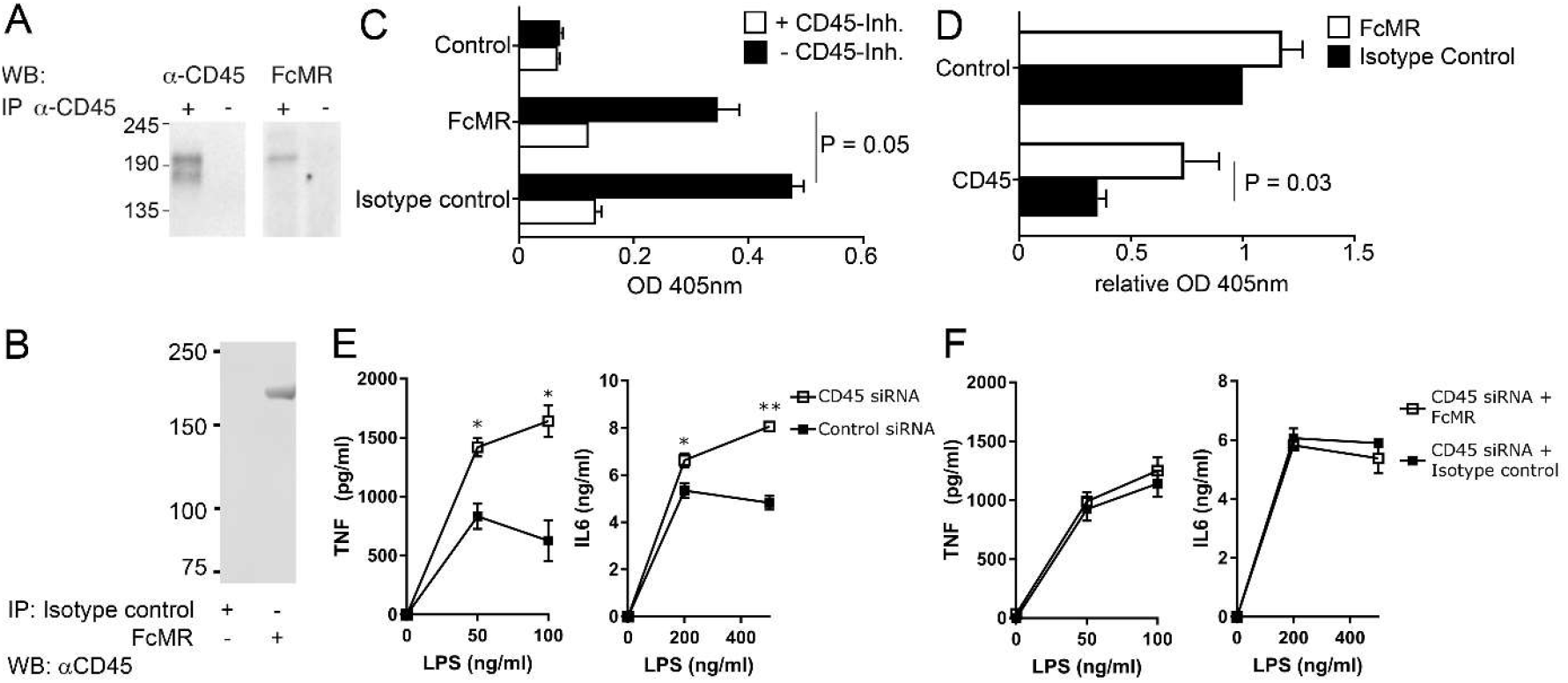
MR inhibits phosphatase activity of CD45 on macrophages. A) Macrophage cell lysates were immune precipitated using a CD45-specific antibody and stained for FcMR binding by far Western Blot. B) Macrophage lysates were immune precipitated with FcMR and stained for CD45 by Western Blot. C) CD45 was precipitated from lysates of FcMR- or isotype control-treated macrophages and incubated with 4-NPP in the presence or absence of the CD45 inhibitor SF1670. Graph depicts CD45-mediated dephosphorylation of 4-NPP measured by colorimetry. Samples without CD45 antibody were used as controls. D) CD45 was precipitated from lysates of FcMR- or isotype control-treated macrophages and incubated with the CD45 substrate TATEGQ-pY-QPQ. Graph depicts the phosphorylation status of TATEGQ-pY-QPQ. Samples without CD45 antibody were used as controls. E) Secretion of TNF and IL-6 by LPS-stimulated macrophages after siRNA-mediated down-regulation of CD45. F) Influence of FcMR on secretion of TNF and IL-6 by LPS-stimulated macrophages after siRNA-mediated down-regulation of CD45. All graphs are depicted as mean ± SEM; for all experiments, n ≥ 3. See also Figure S3.

CD45 can be expressed as different isoforms, depending on alternative splicing of its three exons A, B and C. To identify the CD45 isoform interacting with sMR, we assessed their respective expression using isoform-specific antibodies. Analysis by Western Blot and flow cytometry clearly showed the absence of exons A, B and C in bone marrow-derived macrophages (Figure S3B), pointing out that these cells only express the CD45RO isoform. This is in accordance with our RNA-seq data, which showed a specific read coverage of all exons of *Cd45* except for exons A, B and C (Figure S3C). Additionally, we showed that primary macrophages from spleen, white adipose tissue (WAT), liver and the peritoneal cavity also expressed the CD45RO isoform (Figure S3D), which is in agreement with previous literature (Pilling et al., 2009). Accordingly, we confirmed the direct interaction of FcMR with CD45RO from primary splenic macrophages by far Western Blot (Figure S3E).

Since little is known about CD45 phosphatase activity in macrophages, we next investigated whether CD45 is active in these cells. Therefore, we immunoprecipitated CD45 from macrophage lysates and added 4-nitrophenyl phosphate (pNPP), from which dephosphorylation by CD45 can be quantified using colorimetry. We monitored a clear phosphatase activity, which was blocked by a CD45-specific inhibitor (Figure S3F), demonstrating the presence of active CD45 in macrophages. Next, we tested the effect of sMR on CD45 phosphatase activity. To this end, we immunoprecipitated CD45 from lysates of FcMR-treated macrophage and showed that dephosphorylation of pNPP was reduced when compared to isotype control-treated cells (Figure 3C), indicating that the MR inhibited CD45 phosphatase activity. In a second approach, we assessed the dephosphorylation of a synthetic peptide containing pY505 of Lck, a specific substrate of CD45. We showed that pre-incubation of macrophages with FcMR increased pY505 phosphorylation (Figure 3D), confirming the inhibitory effect of the MR on CD45 phosphatase activity.

To investigate whether sMR-mediated inhibition of CD45 phosphatase activity plays a role in macrophage activation, we down-regulated CD45 expression using siRNA (Figure S3G). Similar to inhibition of CD45 by sMR, CD45 down-regulation resulted in increased expression of TNF and IL-6 after stimulation with LPS (Figure 3E). Importantly, addition of FcMR after down-regulating CD45 had no further effect on cytokine secretion (Figure 3F), demonstrating that the activating effect of the MR on macrophages was indeed due to its inhibition of CD45.

### sMR-mediated inhibition of CD45 activates a Src/Akt/NF-κB signaling cascade in macrophages

We next investigated how sMR-mediated inhibition of CD45 results in macrophage reprogramming towards a pro-inflammatory phenotype. First, we screened for over-represented transcription factor (TF) binding motifs in the non-protein coding regions of FcMR-specific upregulated DE genes. Network visualization of enriched TF binding motifs and their potential target DE genes clearly exposed NF-κB as the dominating transcriptional regulator of differential gene expression across all three time points (Figure 4A). Indeed, macrophage treatment with FcMR significantly downregulated IκBα (Figure 4B), an inhibitor of NF-κB which disables its nuclear translocation, retaining NF-κB in the cytosol. Accordingly, enhanced nuclear translocation of both NF-κB subunits p65 and p50 was observed after FcMR treatment (Figure 4C).

**Figure 4:**
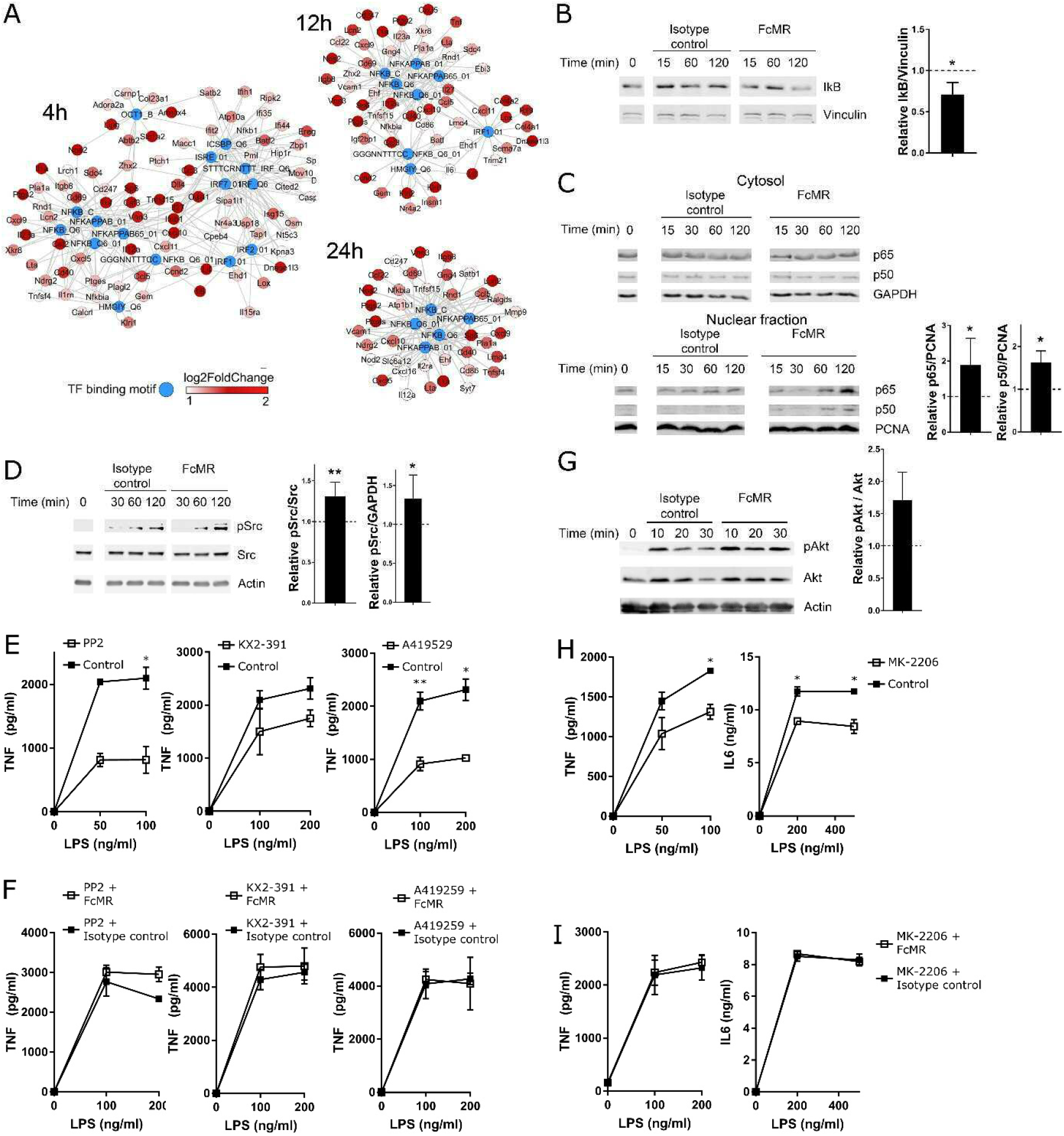
MR reprograms macrophages to a pro-inflammatory phenotype via Src/Akt/NF-κB signaling. A) Network visualization of significantly enriched (q-value < 0.1) TF binding motifs (blue) of the MSigDB motif gene set and their potential targets (colored in red according to their FC) among the upregulated DE genes after 4 h, 12 h or 24 h of FcMR treatment. B) MR-deficient macrophages were treated with FcMR or isotype control for 60 min. Total IκB was determined by Western Blot. C) MR-deficient macrophages were treated with FcMR or isotype control for 60 min. p65 and p50 were monitored in the cytosolic and nuclear fraction by Western Blot. D) MR-deficient macrophages were treated with FcMR or isotype control for 60 min. Src and phosphorylated Src (pSrc) were determined by Western Blot. E) MR-deficient macrophages were treated with 3 μM PP2, 1 μM KX2-391 or 1 μM A419529 and stimulated with LPS. TNF secretion was monitored by ELISA. F) FcMR-treated MR-deficient macrophages were incubated with PP2, KX2-391 or A419259 and stimulated with LPS. Secretion of TNF was determined by ELISA. G) MR-deficient macrophages were treated with FcMR or isotype control for 30 min. Akt and phosphorylated Akt (pAkt) were determined by Western Blot. H) MR-deficient macrophages were treated with 5 μM MK-2206 and stimulated with LPS. TNF secretion was monitored by ELISA. I) FcMR-treated MR-deficient macrophages were incubated with MK-2206 and stimulated with LPS. Secretion of TNF was determined by ELISA. All graphs are depicted as mean ± SEM; for all experiments, n ≥ 3. See also Figure S4.

Subsequently, we aimed at identifying the signaling cascade leading from FcMR-mediated inhibition of CD45 to activation of NF-κB. Since CD45 can lead to the activation of Src kinases (Shrivastava et al., 2004), Src in turn can activate Akt (Chen, 2010) and both Src and Akt have been associated with NF-κB activation (Abu-Amer et al., 1998; Bai et al., 2009; Cheng et al., 2011; Xie et al., 2013), we investigated whether FcMR-mediated inhibition of CD45 resulted in NF-κB activation through signaling via Src and Akt. Indeed, FcMR treatment increased phosphorylation and hence activation of Src (Figure 4D). Furthermore, blocking Src using three different chemical inhibitors (PP2, KX2-391 and A419259) markedly decreased TNF secretion (Figure 4E). Of note, the effect of FcMR on TNF secretion was abolished in the presence of these Src inhibitors (Figure 4F), demonstrating that FcMR-induced macrophage activation depends on Src signaling. Similarly, FcMR treatment clearly increased phosphorylation of Akt (Figure 4G) and addition of an Akt-specific inhibitor decreased LPS-induced secretion of TNF and IL-6 (Figure 4H). Also here, the stimulatory effect of FcMR on TNF secretion was abolished by Akt inhibition (Figure 4I), showing an important role for Akt signaling in FcMR-enhanced TNF secretion. Accordingly, inhibition of Akt prevented FcMR-induced translocation of NF-κB into the nucleus (Figure S4).

Taken together, these data demonstrate that sMR-mediated inhibition of CD45 results in activation of a Src/Akt signaling pathway leading to nuclear translocation of NF-κB and macrophage reprogramming towards an inflammatory phenotype.

### Serum sMR is up-regulated in obesity and promotes HFD-induced metabolic dysfunctions and hepatic steatosis

Next, we monitored whether the inflammatory effect of the MR on macrophages regulates inflammatory processes *in vivo* using a murine model of obesity-induced metaflammation. We first investigated whether high fat diet (HFD) feeding resulted in changes in serum sMR levels (Figure S5A) and we demonstrated significantly increased sMR concentrations in the serum of HFD-fed obese mice, as compared to lean control diet (CD)-fed mice (Figure 5A). Additionally, serum sMR levels positively correlated with body weight and fat mass of the mice (Figures 5B, 5C). In humans, serum sMR levels were also increased in obese individuals when compared to lean subjects (Figure 5D), and correlated positively with body mass index (BMI) and fat mass (Figures 5E 5F), indicating a direct correlation between serum sMR levels and obesity in both humans and mice.

**Figure 5:**
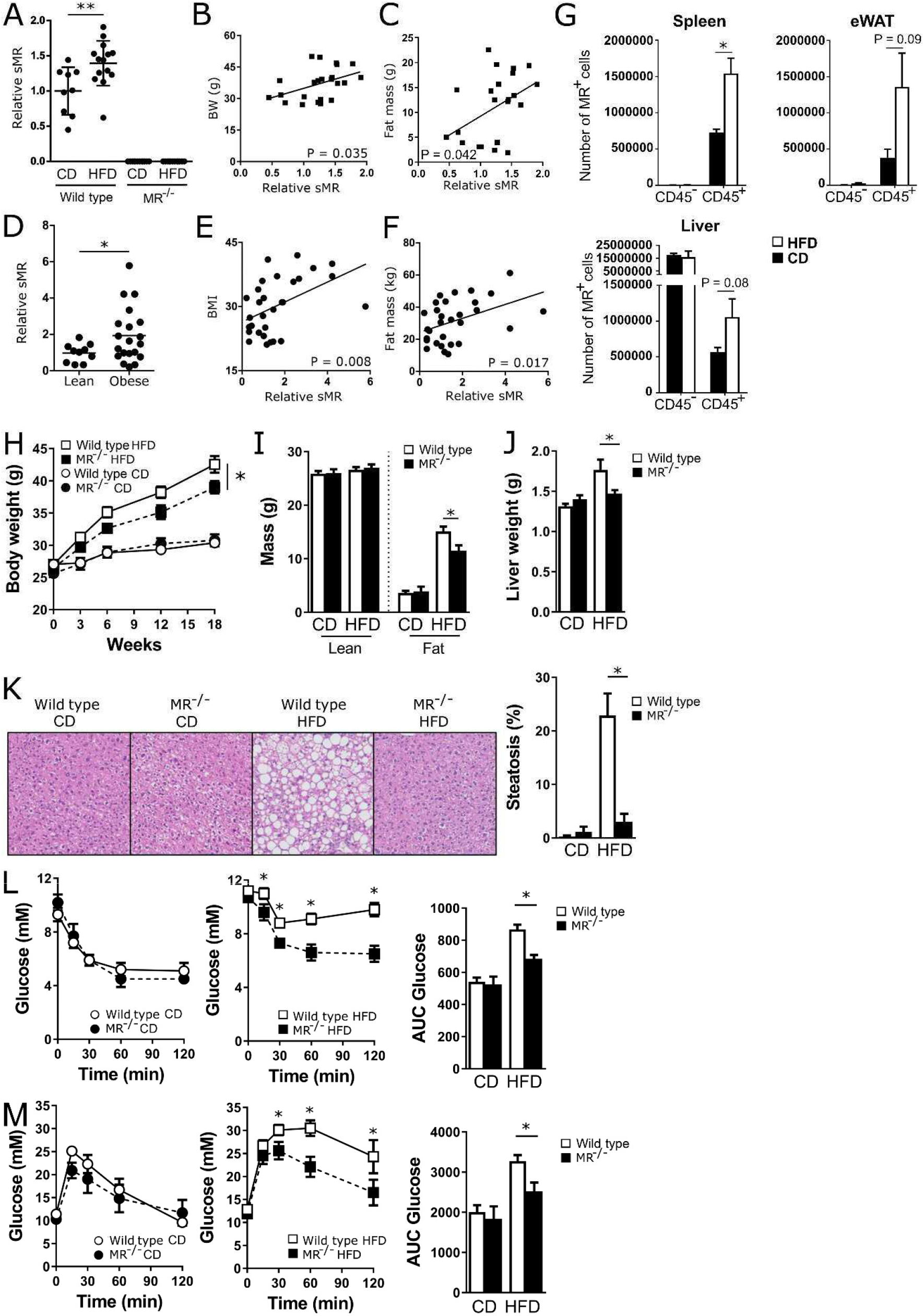
sMR is up-regulated in obesity and regulates HFD-induced metabolic dysfunctions and hepatic steatosis. A) sMR levels in the serum of wild-type or MR-deficient mice after HFD or CD feeding. B-C) Correlation of sMR serum levels and body weight (B) or fat mass (C) of all mice depicted in A). D) sMR levels in the serum of lean and obese humans. E-F) Correlation between sMR serum levels and BMI (E) or fat mass (F). G) MR expression in different organs after HFD or CD feeding. H) Body weight of mice on HFD or CD for 18 weeks. I) Lean and fat mass after 18 weeks of diet determined by MRI. J) Liver weight after 18 weeks of diet. K) H&E staining to monitor steatosis. L) Intraperitoneal insulin tolerance test. Blood glucose levels were measured at the indicated time-points and the AUC of the glucose excursion curve was calculated as a surrogate measure for whole-body insulin resistance. M) Intraperitoneal glucose tolerance test. The AUC of the glucose excursion curve was calculated as a surrogate measure for whole-body glucose tolerance. eWAT, epidydimal white adipose tissue. mWAT, mesenteric WAT. iWAT, inguinal WAT. BAT, intrascapular brown adipose tissue. Results are expressed as means ± SEM; n=3 mice per group for G, n=10-15 mice per group for H-K; n= 5-10 mice per group for L-M. See also Figure S5.

Subsequently, we analyzed changes in MR expressing cells as putative source for increased sMR serum levels in HFD-fed mice. In spleen and white adipose tissue (WAT) of both CD- and HFD-fed mice, nearly all MR-expressing cells were CD45^+^, whereas in liver, CD45^-^ cells also expressed the MR. These latter cells were identified as CD31^+^CD146^+^ liver sinusoidal endothelial cells (LSECs), which were indeed previously reported to express the MR (Martinez-Pomares, 2012). Importantly, whereas no differences in MR expression could be detected in CD45^-^ cells, a clear increase in MR^+^ cells was observed in CD45^+^ hematopoietic cells in spleen, liver and WAT of HFD-fed obese mice compared to CD-fed mice (Figure 5G). Of note, CD45^+^MR^+^ cells in all three organs were mainly identified as CD64^+^F4/80^+^ macrophages (Figure S5B). Taken together, this demonstrates that obesity increased MR-expressing macrophages in spleen, liver and WAT.

To test whether increased sMR levels regulate macrophage-mediated inflammatory diseases *in vivo*, we then analyzed the development of obesity-induced metaflammation in MR-deficient mice. Whereas no differences in body weights were found between wild-type and MR-deficient mice on CD, MR-deficient mice were partially protected from weight gain on a HFD (Figure 5H). Analysis of body composition showed that the lower body weight in HFD-fed MR-deficient mice resulted exclusively from a reduction in fat mass, without affecting lean mass (Figure 5I). Accordingly, the weights of epididymal, mesenteric and subcutaneous (inguinal) WAT, as well as brown adipose tissue (BAT), were significantly lower in MR-deficient mice on HFD (Figure S5C). In addition, liver weight was also markedly lower in HFD-fed MR-deficient mice as compared to wild-type controls (Figure 5J), suggesting a reduction in hepatic steatosis. Indeed, MR-deficient mice were completely protected against HFD-induced hepatic steatosis (Figure 5K). Accordingly, hepatic triglycerides, total cholesterol and phospholipids contents (Figure S5D) and circulating alanine aminotransaminase (ALAT) levels (Figure S5E) were markedly decreased in HFD-fed MR-deficient mice.

HFD-fed MR-deficient mice displayed significantly lower fasting plasma insulin levels than wild-type mice, whereas fasting glucose levels were unchanged (Figure S5F). The calculated HOmeostasis Model Assessment of Insulin Resistance index (HOMA-IR) adjusted for mice was significantly reduced in HFD-fed MR-deficient mice (Figure S5G), suggesting that insulin resistance is less severe in these mice. In line with this, whole-body insulin sensitivity (Figure 5L) and glucose tolerance (Figure 5M) were significantly higher in HFD-fed MR-deficient mice compared to wild-type mice. Of note, the alleviated hepatic steatosis and whole-body metabolic homeostasis were still observed when HFD-fed MR-deficient mice were weight-paired to their wild-type counterparts (Figure S5H), indicating that MR deficiency protects against HFD-induced metabolic dysfunctions independent of body weight changes. Altogether, these data demonstrate that the MR contributes to obesity-induced metabolic dysfunctions.

### MR promotes inflammatory macrophage accumulation in eWAT and liver during obesity

Since our *in vitro* data demonstrate that the MR reprograms macrophages towards an inflammatory phenotype, we next investigated whether the observed metabolic changes in MR-deficient mice might be caused by reduced pro-inflammatory macrophage activation in metabolic tissues.

As previously reported, HFD significantly increased obesity-associated pro-inflammatory CD11c^+^ adipose tissue macrophages (ATMs) in eWAT of wild-type mice (Lumeng et al., 2007), whereas total ATMs and CD11b^+^Ly6C^+^ monocytes were not affected (Figures 6A, S6A, S6B). Remarkably, while no significant differences in total ATMs and monocyte numbers were observed between genotypes, inflammatory CD11c^+^ ATM numbers were found to be significantly higher in HFD-fed MR-expressing wild-type mice as compared to MR-deficient mice (Figure 6A).

**Figure 6:**
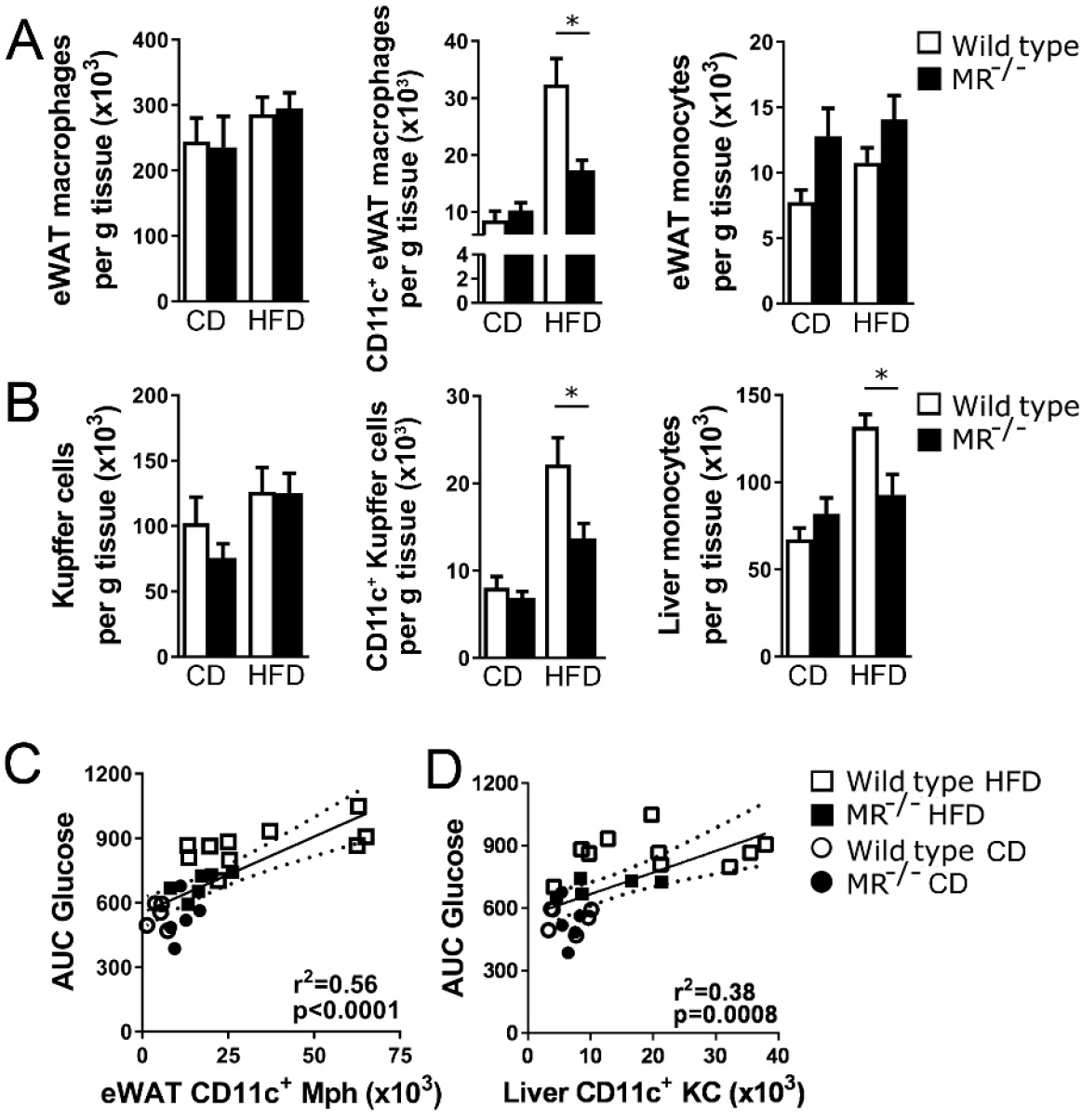
MR regulates WAT and liver macrophage activation after HFD feeding. A) Numbers of total macrophages, CD11c^+^ macrophages and monocytes per gram eWAT in HFD- or CD-fed wild-type or MR-deficient mice determined by flow cytometry. B) Numbers of total macrophages, CD11c^+^ macrophages and monocytes per gram liver in HFD- or CD-fed wild-type or MR-deficient mice determined by flow cytometry. C-D) Correlation between CD11c^+^ macrophages and whole-body insulin sensitivity, assessed by the area under the curve (AUC) of the glucose excursion curve in eWAT (C) and liver (D). Results are expressed as means ± SEM; n=10-15 mice per group. See also Figure S6.

In the liver, HFD significantly increased pro-inflammatory CD11c^+^ KCs and CD11b^+^Ly6C^+^ monocytes in wild-type mice, while total KCs were not affected (Figure 6B). Similar to what was observed in eWAT, inflammatory CD11c^+^ KCs, but also monocytes, were more abundant in liver of MR-expressing wild-type mice as compared to MR-deficient mice, while total KCs were not affected (Figure 6B). This was associated with higher expression of genes involved in pro-inflammatory macrophage activation in liver and WAT of MR-expressing wild-type mice (Supplementary Figure S6C). The differences in pro-inflammatory macrophage abundances in metabolic tissues were still present when wild-type and MR-deficient mice were weight-paired (Figures S6D, S6E), demonstrating that also the regulation of obesity-induced pro-inflammatory macrophages by the MR is independent of changes in body weight. Additionally, a strong positive correlation between the numbers of pro-inflammatory macrophages in eWAT and liver, and whole-body insulin resistance was observed (Figures 6C, 6D).

### sMR treatment induces pro-inflammatory cytokines, metabolic dysfunctions and increased pro-inflammatory macrophages

Finally, we investigated whether *in vivo* administration of sMR in lean mice is able to induce macrophage activation and metabolic dysfunctions. Therefore, we first monitored circulating cytokine levels in response to a single intraperitoneal injection of FcMR in CD-fed mice. In accordance with our *in vitro* experiments, even a single injection of FcMR acutely increased serum levels of TNF and IL-6 and of the chemokine MCP-1/CCL2 compared to isotype control-treated mice (Figure 7A).

**Figure 7:**
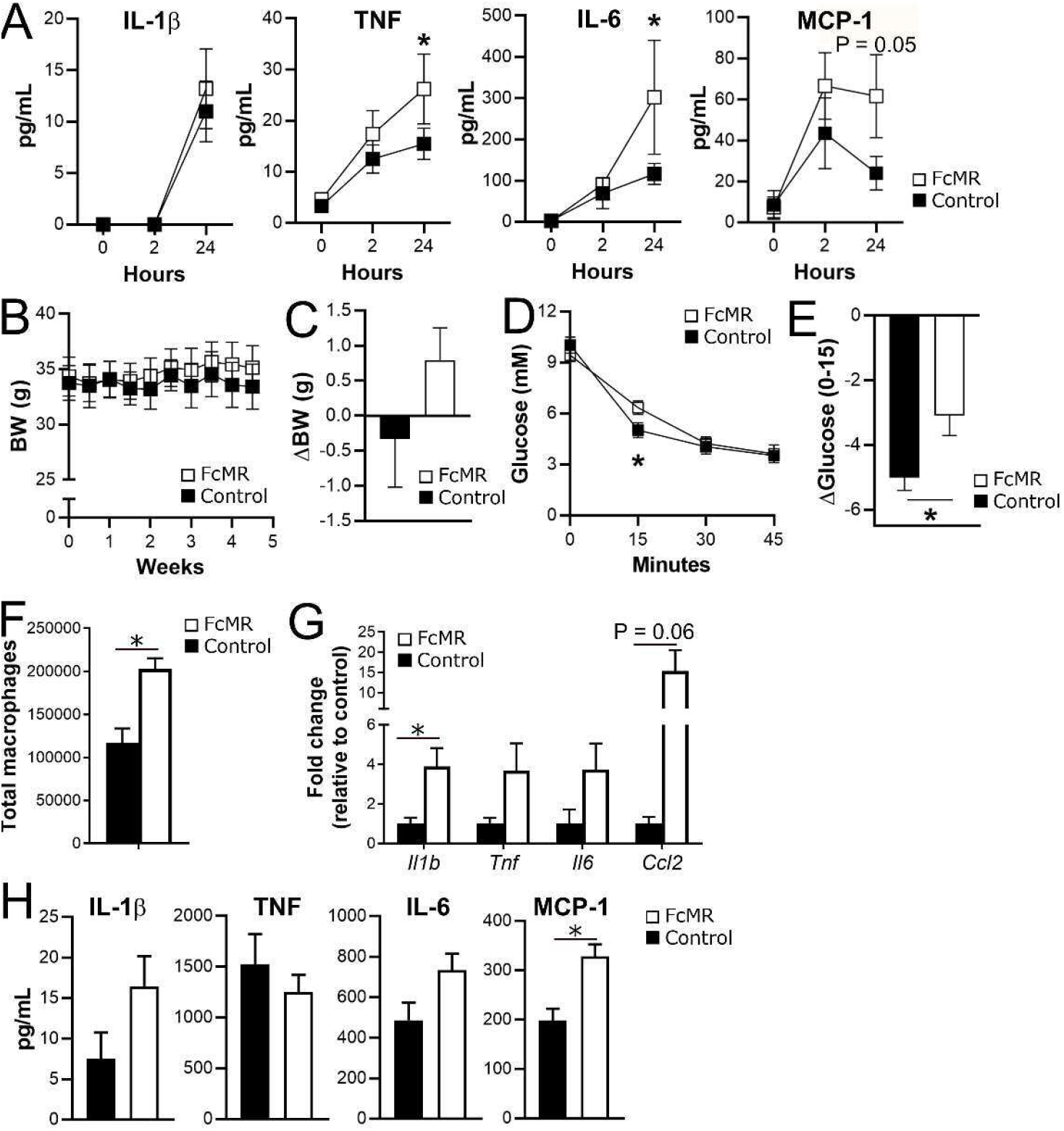
Increased MR levels regulate whole-body metabolism and promote inflammation. A) CD-fed wild type mice were injected i.p. with 4,82 μmoles/mouse FcMR or isotype control. Serum cytokine concentrations were determined by CBA at the indicated timepoints. B-C) Wild-type mice were fed a CD and concomitantly injected i.p. with 4,82 μmoles/mouse FcMR or isotype control every 3 days. Graphs depict body weight over time (B) and overall change in body weight (C). D-E) Intraperitoneal insulin tolerance test after FcMR treatment of CD-fed mice. Graph in E) shows changes in circulating Glucose levels at 15 min post insulin i.p. F) The number of F4/80^+^ MACS sorted eWAT macrophages was determined after 4 weeks of treatment. G) Expression of inflammatory genes in eWAT was monitored by qPCR. H) Cytokine secretion by F4/80^+^ macrophages from eWAT was determined by CBA after stimulation with 100 ng/ml LPS. BW: body weight. Results are expressed as means ± SEM; n=4-5 mice per group.

To assess whether increased sMR levels also promote whole-body metabolic dysfunctions, we injected lean mice with FcMR or isotype control every three days. After four weeks of treatment, we monitored a mild increase in body weight in FcMR-treated mice compared to control mice (Figures 7B, 7C). In addition, insulin sensitivity, as measured by an acute drop in blood glucose levels following insulin i.p. injection, was reduced in FcMR-treated mice compared to control mice (Figures 7D, 7E), confirming the detrimental effect of the sMR on whole-body metabolic homeostasis. Of note, when mice were fed a HFD concomitantly with FcMR treatment for four weeks, the effect of FcMR on HFD-induced insulin resistance was even more pronounced (Figures S7A, S7B, S7C).

FcMR treatment increased macrophage numbers in eWAT of lean mice (Figure 7F). Moreover, gene expression of *Il1b, Tnf, Il6* and *Ccl2* was increased in eWAT of FcMR-treated lean mice (Figure 7G). Accordingly, macrophages isolated from these mice showed increased secretion of most of these cytokines upon stimulation with LPS (Figure 7H), demonstrating that, in lean mice, increased serum sMR levels induce the secretion of pro-inflammatory cytokines, induce whole-body insulin resistance and promote macrophage activation in metabolic tissues *in vivo*.

## Discussion

The MR as a member of the C-type lectin family has mainly been described as an endocytic receptor recognizing glycosylated antigens and mediating antigen processing and presentation (Rauen et al., 2014; Kreer et al., 2017). However, the extracellular region of the MR can be shed by metalloproteases and released as a soluble protein in the extracellular space. Consequently, the sMR is detectable in murine and human serum (Martínez-Pomares et al., 1998; Jordens et al., 1999), and recent studies reported an increase of serum sMR levels in a variety of inflammatory diseases and serum sMR levels directly correlated with severity of disease and mortality (Andersen et al., 2014; Rødgaard-Hansen et al., 2015; Ding et al., 2017; Suzuki et al., 2018; Loonen et al., 2019; Saha et al., 2019) Here, we demonstrated that, in addition to a mere phenotypic correlation, the sMR plays a direct, functional role in macrophage activation, driving reprogramming towards a pro-inflammatory phenotype. By interacting with and inactivating CD45, the sMR reprograms macrophages via activation of Src/Akt signaling and nuclear translocation of NF-κB. *In vivo,* sMR levels were increased in obese mice and humans as compared to lean controls, and we found that MR deficiency reduced adipose tissue and liver pro-inflammatory macrophages and protected against obesity-induced metabolic dysfunctions. Consistently, treatment of lean mice with sMR acutely increased serum pro-inflammatory cytokines, and induced both tissue macrophage activation and systemic insulin resistance.

In this study, we found increased MR^+^ cells in spleen, liver and eWAT upon HFD feeding, which were almost exclusively macrophages. Since cell surface MR can be shed and released as a soluble form, obesity-induced changes in tissue homeostasis may increase MR expression and sMR release by macrophages, creating increased local and systemic sMR levels to promote macrophage-mediated inflammation and metabolic dysfunctions. An important factor in this process could be the ligand-inducible transcription factor PPARγ, which is activated, among others, by free fatty acids. Indeed, single cell RNA sequencing analysis of adipose tissue immune cells revealed that PPAR signaling is among the upregulated pathways in obesity-induced lipid-associated macrophages (LAMs) in both mice and humans (Jaitin et al., 2019). Since *Mrc1* (encoding the MR) is also a direct target gene of PPARγ (Klotz et al., 2009), obesity-induced activation of PPARγ in macrophages may lead to enhanced transcription of the MR, eventually resulting in increased sMR levels.

Our data demonstrate that macrophage activation by the MR was due to MR-mediated inhibition of CD45, which in turn leads to activation of Src and Akt, and nuclear translocation of NF-κB. CD45 has been postulated to inhibit Src kinases (Shrivastava et al., 2004) and Akt (Chen, 2010), a known regulator of NF-κB (Abu-Amer et al., 1998; Lluis et al., 2007; Xie et al., 2013). Here, we show that CD45-mediated dephosphorylation of Src induces Akt-mediated nuclear translocation of NF-κB, and that the sMR uses this novel signaling pathway to induce macrophage reprogramming towards an inflammatory phenotype. Of note, it has been reported that activation of Src kinases can lead to activation of NF-κB independent of Akt. Such signaling has been demonstrated to be mediated by the Bruton’s tyrosine kinase (BTK) (Doyle et al., 2005; Weber et al., 2017). Here, MR-induced macrophage activation was not affected by BTK, as inhibition of BTK by Ibrutinib did not influence secretion of TNF and IL-6 by macrophages (data not shown). In contrast, MR-mediated macrophage activation depended on Akt signaling, as its inhibition abrogated MR-mediated nuclear translocation of NF-κB and ensuing TNF and IL-6 secretion by macrophages. Additionally, Akt has been postulated as a regulator that can fine tune NF-κB-mediated responses through regulating efficient binding of p65 to specific target promoters (Cheng et al., 2011). Of note, these authors demonstrated that NF-κB-mediated expression of TNF was particularly sensitive to Akt signaling, which is in accordance with the Akt-dependency of TNF expression after MR-induced activation of macrophages described here.

The immunometabolic phenotype of obese MR-deficient mice resembles that of mice deficient for MGL/CLEC10A, another member of the C-type lectin family. Indeed, these mice displayed reduced hepatic steatosis, insulin resistance and glucose intolerance upon HFD when compared to wild-type mice, a feature that was associated with lower AT pro-inflammatory macrophages (Westcott et al., 2009). In another context, MGL/CLEC10A was also shown to bind and inactivate CD45 (van Vliet et al., 2006), offering the possibility that MGL/CLEC10A can directly induce macrophage reprogramming by inhibition of CD45, similar to the MR. Future studies will have to reveal whether MGL/CLEC10A indeed plays a direct role in macrophage activation, whether its expression is also increased in HFD-induced obesity, and whether this may be mediated by a soluble form of MGL/CLEC10A.

In summary, we demonstrate that a soluble form of the MR reprograms macrophages towards an inflammatory phenotype by interacting with CD45 on the surface of these macrophages. MR-mediated inhibition of CD45 activated Src and Akt kinases, leading to nuclear translocation of NF-κB and induction of a transcriptional program that ultimately results in enhanced inflammatory cytokine production. Furthermore, sMR levels in serum of obese mice and humans are increased, strongly correlating with body weight and adiposity. Accordingly, MR deficiency resulted in fewer adipose tissue and liver pro-inflammatory macrophages and protection against hepatic steatosis and metaflammation, whereas increased MR levels induced elevated serum pro-inflammatory cytokines, macrophage activation and metabolic dysfunctions. Altogether, our results identify sMR as a novel regulator of pro-inflammatory macrophage activation and could contribute to the development of new therapeutic strategies for metaflammation and other hyperinflammatory diseases. Targeting MR-mediated activation of macrophages using antibodies, nanobodies, aptamers or small molecules that could prevent MR interacting with macrophage CD45RO might open new possibilities for therapeutics aimed at dampening (meta-)inflammation.

## Acknowledgements

This work is funded by the Deutsche Forschungsgemeinschaft (DFG, German Research Foundation) under Germany’s Excellence Strategy – EXC2151 – 390873048 (to SB), an EFSD/Lilly Research Grant Fellowship from the European Federation for the Study of Diabetes (to BG), the NWO Graduate School Program 022.006.010 (to HJPvdZ), and the Dutch Organization for Scientific Research (ZonMW TOP Grant 91214131, to BG and MY). We thank Frank Otto and Arifa Ozir-Fazalalikhan for their precious technical help.

The authors declare no competing interests.

## Author contributions

Conceptualization: B.G. and S.B.; Methodology: B.G., S.B., J.L.S., M.E., H.J.P.vdZ, L.H., K.H., J.S.-S., V.vH, B.E.; Investigation: B.G., S.B., M.E., H.J.P.vdZ, L.H., J.S.-S., L.S., I.S., N.G-T, J.L., A.Z, L.H., K.dR, M.W, V.vH, L.R.P., B.E.; Writing – Original Draft: B.G, S.B., H.J.P.vdZ, M.E, J. S.-S., J.L.S.; Writing – Review & Editing, H.J.P.vdZ, S.B., B.G; Funding Acquisition:, S.B., B.G, J.L.S., M.Y., H.J.P.vdZ; Resources: M.W, H.P., K.WvD; Supervision: S.B. B.G., J.L.S., K.H., M.Y., V.vH, K.WvD, B.E.

## STAR Methods

### Key resources table

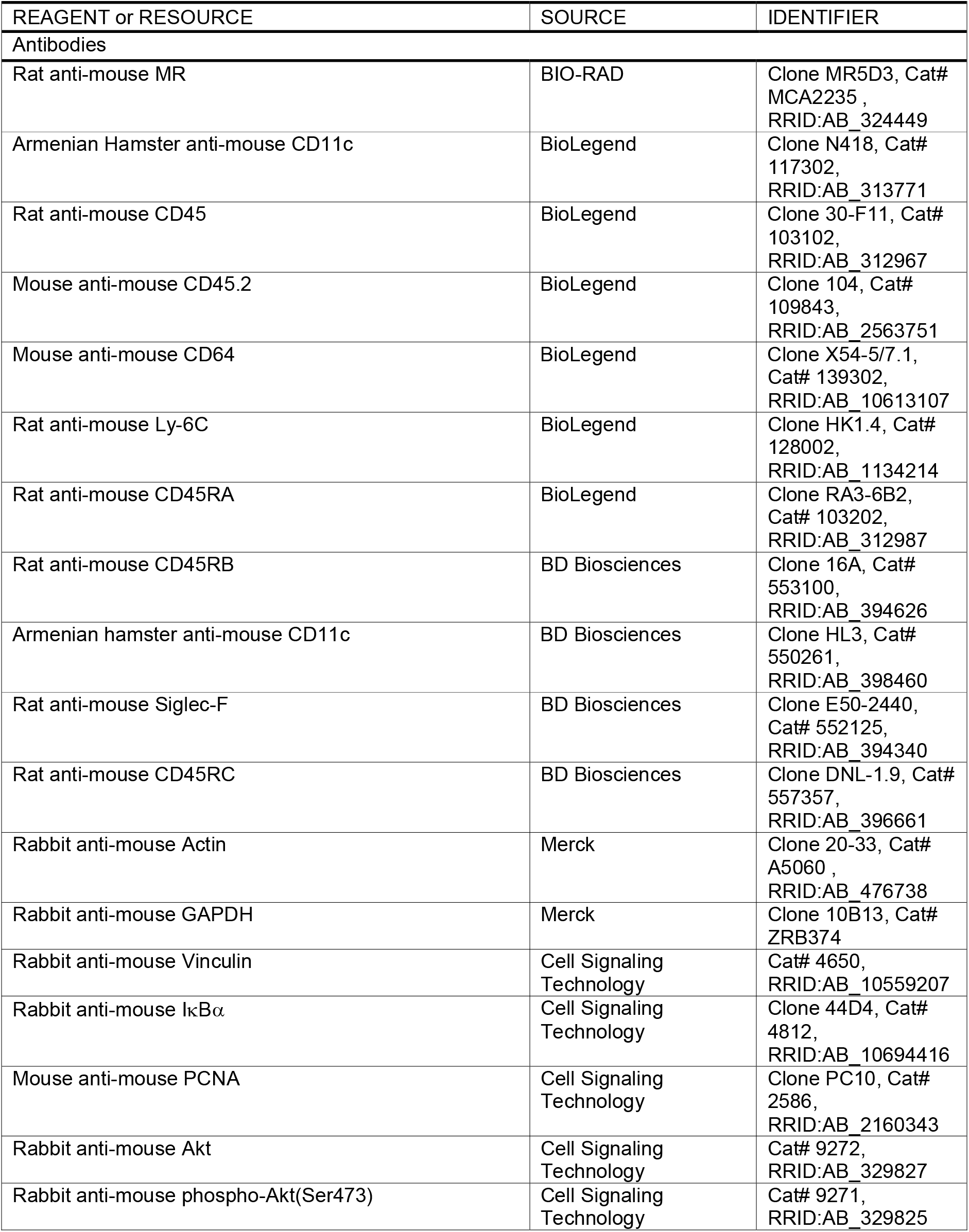
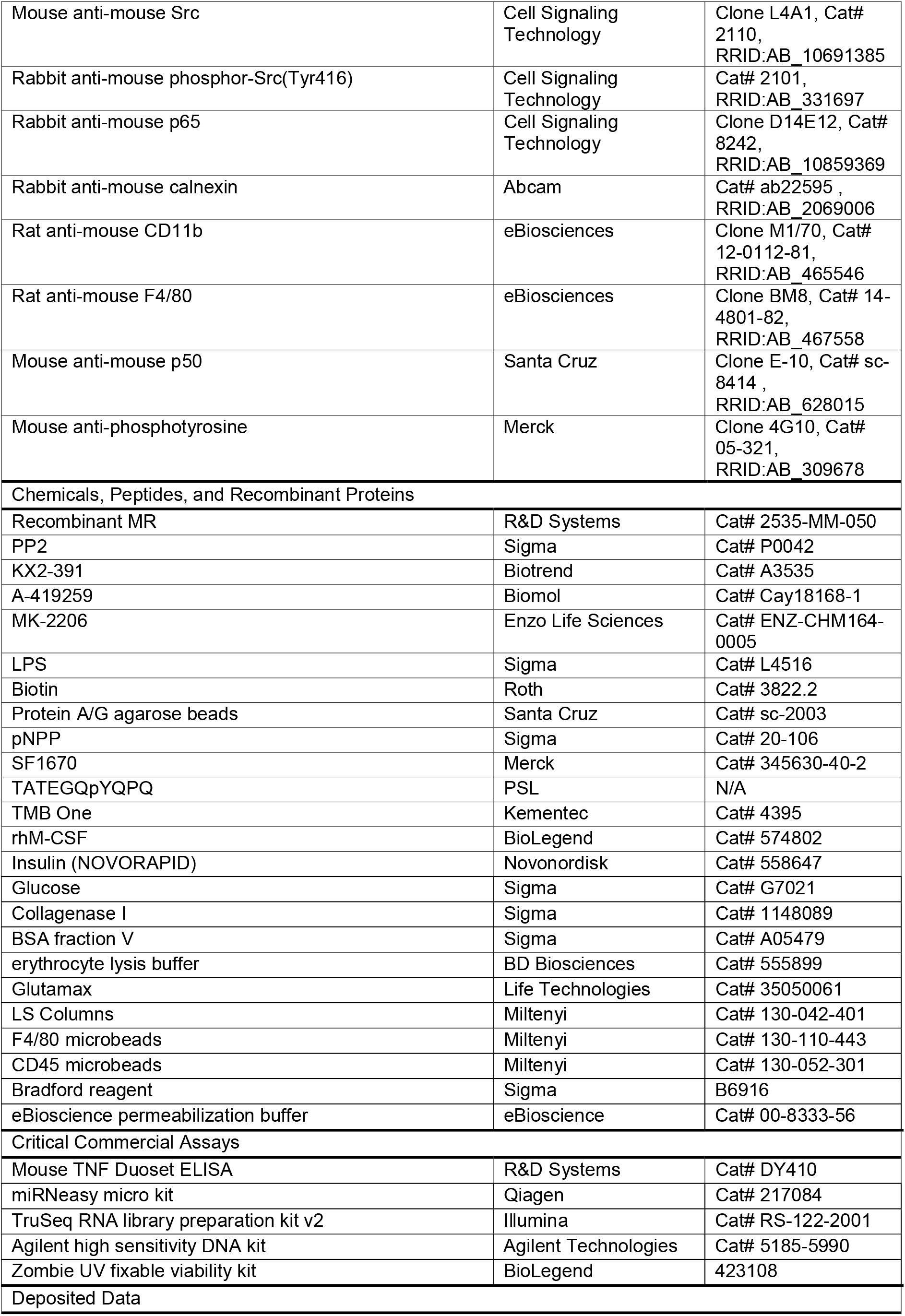
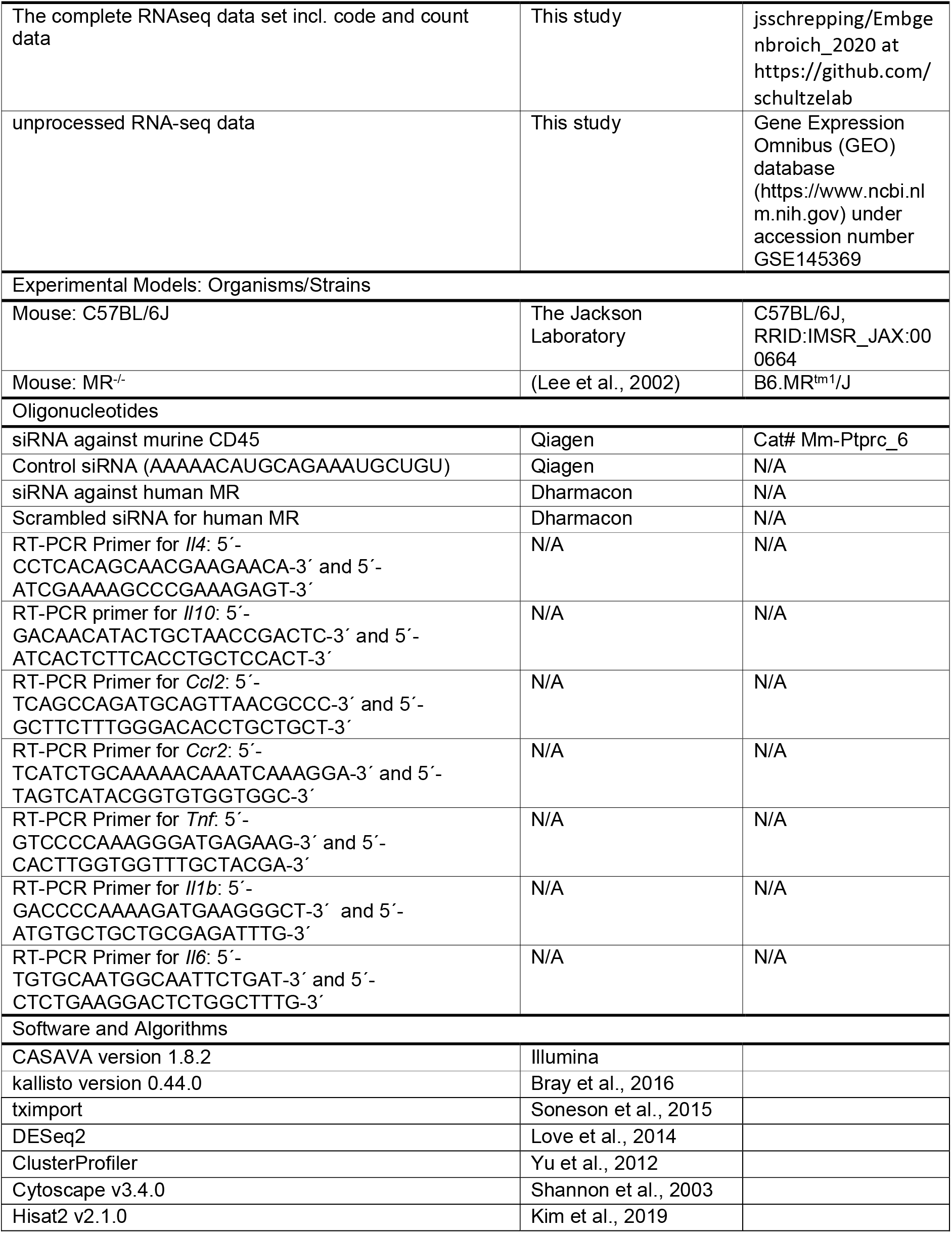

### Contact for reagent and resource sharing

Further information and requests for resources and reagents should be directed to and will be fulfilled by the Lead Contact, Sven Burgdorf

### Experimental Model and subject details

#### Generation of bone marrow-derived macrophages

Bone marrow was flushed from the femurs and tibias of mice and cultured for 7 days in medium containing 2.5 % supernatant of a GM-CSF-producing cell line (total concentration 150 ng/ml).

#### Mice and diet

All animal experiments were performed in accordance with the Guide for the Care and Use of Laboratory Animals of the Institute for Laboratory Animal Research and have received approval from the university Ethical Review Boards (DEC No. 12199; Leiden University Medical Center, Leiden, The Netherlands). Male C57BL/6 MR^-/-^ mice and age-matched C57BL/6 wild-type mice from the same breeding facility were housed in a temperature-controlled room with a 12 hour light-dark cycle. Throughout the experiment, food and tap water were available ad libitum. 8-10 weeks old male mice were randomized according to total body weight, lean and fat mass, and fasting plasma glucose, insulin, TC and TG levels, after which they were fed a high-fat diet (HFD, 45% energy derived from fat, D12451, Research Diets) or a control diet (CD, 10% energy derived from fat, D12450B, Research Diets) for 18 weeks.

For *in vivo* FcMR treatment, C57BL/6 wild-type mice were randomized as above at baseline or after a run-in HFD of 14 weeks. Subsequently, mice were biweekly intraperitoneally injected with 50 μg FcMR or 6.75 μg isotype control, to yield the same administered dose of hIgG1, for four weeks while maintaining the HFD.

#### Human samples

Serum samples from twenty-six healthy, weight-stable, nonsmoking Caucasian volunteer subjects, 12 lean (2 males, 10 females, BMI 23.3 +/− 0.5 kg/m2) and 14 obese (2 males, 12 females, BMI 35.2 -/- 1.2 kg/m2), this latter before and after weight loss, were collected in the framework of a clinical trial (Wijngaarden et al., 2013) and used to measure circulating sMR in a subset of still available samples. This study (Clinical Trial Registration No. NTR2401) was approved by the Medical Ethics Committee of the Leiden University Medical Centre and performed in accordance with the principles of the revised Declaration of Helsinki. All volunteers gave written informed consent before participation.

### Method details

#### Generation and purification of FcMR

FcMR proteins (encompassing the CR region, the FN II domain and CTLD1-2 fused to the Fc region of hIgG1) and isotype controls (Fc region of hIgG1) were generated as described previously (Martinez-Pomares et al., 2006). For all *in vitro* experiments, FcMR and isotype controls were used in a concentration of 10 μg/ml.

#### Purification of sMR from the supernatant of MR-expressing cells

Supernatant of bone marrow-derived macrophages was collected and loaded on an affinity chromatography column containing Sepharose beads that were covalently linked to an anti-MR antibody (MR5D3, BIO-RAD). After extensive washing, sMR was eluted in 0.1 M Glycin (pH 2.5), neutralized with 1 M Tris (pH 9.0) and dialysed against PBS containing 10% PEG for 24 h.

#### Monitoring secretion of TNF and IL-6

Macrophages were incubated with 10 μg/ml FcMR or isotype control, 300 ng/ml recombinant MR (2535-MM-050, R&D Systems), 30 ng/ml purified sMR, 3 μM PP2, 1 μM KX2-391, 1 μM A419259 or 5 μM MK-2206. After 2 h, LPS was added in the given concentrations. Unless indicated differently, supernatants were collected at 3 h (TNF) or 18 h (IL-6) post LPS stimulation, and secreted cytokine levels were measured by ELISA.

#### Sample preparation for Western Blot analysis

For whole cell lysates, samples were lysed in 10 mM triethanolamine, 150 mM NaCl, 1 mM MgCl2, 1 mM CaCl2 and 1% Triton X-100. For the extraction of nuclear extracts, cells were lysed first in 50 mM HEPES-KOH, 1 mM EDTA (pH 8.0), 140 mM NaCl, 0.25% Triton X100, 0.5% Igepal and 10% glycerol and the cytosolic fraction was harvested. Afterwards, pellets were resuspended in 10 mM Tris-HCl (pH 8.0), 1 mM EDTA, 100 mM NaCl, 0.5 mM EGTA, 0.1% Sodium desoxycholic acid and 0.5% sodium N-lauryl sarcosine, sonicated and centrifuged, yielding the nuclear fraction.

#### Surface biotinylation and co-immuno precipitation experiments

Bone marrow-derived macrophages were incubated with 0.5 mg/ml biotin for 30 min and washed extensively. Afterwards, cells were lysed and 10 μg/ml FcMR was added for 1 h on ice. Subsequently, FcMR was immunoprecipitated using protein A/G-based affinity chromatography and samples were loaded on an SDS-PAGE for analysis by Western Blot using streptavidin or a CD45-specific antibody. Alternatively, a CD45-specific antibody was added to macrophage lysates and precipitated by protein A/G-based affinity chromatography for subsequent far Western Blot analysis using FcMR.

#### CD45 phosphatase assay

CD45 was immunoprecipitated from macrophage lysates and incubated with 2 mM pNPP for 18 h at 37 °C in the presence or absence of 1 μM of the CD45-specific inhibitor SF1670. Dephosphorylation of pNPP was quantified by colorimetry at 405 nm. Alternatively, immunoprecipitated CD45 was incubated with 0.25 μg of the biotinylated peptide TATEGQpYQPQ for 18 h at 37 °C in the presence or absence of SF1670. Phosphorylated TATEGQpYQPQ was monitored after affinity chromatography using streptavidin-agarose, staining with the phosphospecific primary antibody 4G10 (Milipore), a HRP-conjugated secondary antibody and addition of the HRP substrate TMB One (Kementec).

#### siRNA-mediated down-regulation of CD45

siRNA against CD45 (Mm-Ptprc_6 Flexitube siRNA, Qiagen) or control siRNA (AAAAACAUGCAGAAAUGCUGU; containing a specific sequence of the luciferase gene) were obtained from Qiagen. After five days of culture in GM-CSF-containing medium, cells were electroporated with 4 μg siRNA using a Gene Pulser Xcell Electroporation Systems (Bio-Rad) with two sequential pulses of 1000 V for 0.5 msec each. Cells were incubated for 2 days before subsequent experiments were performed.

#### Blood monocyte-derived macrophages and siRNA-mediated down-regulation of MR expression

Human CD14^+^ monocytes were isolated from blood of anonymous healthy volunteers, as described previously (Hussaarts et al., 2013), and cultured in RPMI 1640 (Invitrogen) supplemented with 10% heat-inactivated FSC, 100 U/ml penicillin, 100 μg/ml streptomycin and 50 ng/mL of recombinant human M-CSF (BioLegend) in plates with NunclonTM Delta Surface coating (Nunc). On day 4 of differentiation, cells were electroporated with either 455 nM anti-*Mrc1* siRNA or 455 nM scrambled siRNA (Dharmacon) using the Neon® transfection system (Invitrogen) using one pulse of 1600V for 20 ms. Cells were incubated for 2 days and next incubated for 24 h with 100 ng/ml LPS and 50 ng/ml IFN-γ. Supernatant was harvested after 24 h for analyses of TNF by ELISA using a commercially available kit (BioLegend).

#### RNA isolation and RNAseq analysis

RNA of 5×10^6^ bone-marrow derived macrophages treated with FcMR or isotype control for 4 h, 12 h and 24 h was isolated with trizol and miRNeasy micro kit (Qiagen) according to the manufacturer’s protocol. RNA quality was assessed by visualization of 28S and 18S band integrity on a Tapestation 2200 (Agilent). 100 ng of RNA was converted into cDNA libraries using the TruSeq RNA library preparation kit v2. Size distribution of cDNA libraries was measured using the Agilent high sensitivity DNA assay on a Tapestation 2200 (Agilent). cDNA libraries were quantified using KAPA Library Quantification Kits (Kapa Biosystems). After cluster generation on a cBot, 75 bp single read sequencing was performed on a HiSeq1500.

#### Bioinformatic analysis

After base calling and de-multiplexing using CASAVA version 1.8.2 and subsequent quality control using fastQC, the 75 bp single-end reads were pseudoaligned to the mm10-based mouse Gencode reference transcriptome vM16 using kallisto version 0.44.0 (Bray et al., 2016). Transcript abundance estimations were imported to R and summarized on gene level using tximport (Soneson et al., 2015). Downstream analyses were performed using DESeq2 (Love et al., 2014). After filtering of lowly expressed genes (rowSums > 10) and variance stabilizing transformation, principle component analysis was performed on all present genes using the prcomp package. Differential expression analysis was performed comparing FcMR-treated samples versus controls for each time point without pre-defined log2 fold change threshold and using independent hypothesis weighting (IHW) (Ignatiadis et al., 2016) as the multiple testing procedure. Genes with an adjusted p-value < 0.05 and a fold change (FC) > 2 were determined as significantly differentially expressed. Normalized and z-scaled expression values of the union of differentially expressed (DE) genes over all three time points were visualized in a heatmap. Gene ontology enrichment analyses were performed on those genes shared between all three DE gene sets (shared), as well as the respective gene sets for each time point (4 h, 12 h and 24 h) and those genes unique for each time point (4h.u, 12 h.u and 24 h.u) using the R package ClusterProfiler (Yu et al., 2012) and visualized in a dot plot. Based on the differential expression analysis, genes with significant upregulation in at least two consecutive time points were selected and ranked according to their FC at each timepoint. Normalized and z-scaled expression values of the union of the top 25 genes for each comparison were visualized in a heatmap. Furthermore, enrichment analysis on the FcMR-specific DE genes for each time point was performed based on 49 previously defined, stimulus-specific macrophage expression signatures encompassing 28 different immunological stimuli (Xue et al., 2014), using ClusterProfiler’s “enricher” function. Significantly enriched signatures were visualized in a dot plot and normalized and z-scaled expression values of the genes to the enriched signatures were plotted in a heatmap. To identify transcriptional regulators responsible for the FcMR-induced changes in gene expression, transcription factor motif enrichment analyses was performed using ClusterProfiler’s “enricher” function based on the MSigDB (Liberzon et al., 2011) motif gene sets on the FcMR-specific DE genes for each time point. Motifs with q-value < 0.1 were selected and results were visualized in networks showing the enriched TF motifs and their potential targets among the DE genes of the respective comparison using Cytoscape v3.4.0 (Shannon et al., 2003). For determination of the *Cd45* transcript variant expressed in the cells of this data set, reads were aligned to the reference genome mm10 using Hisat2 v2.1.0 (Kim et al., 2019) and visualized using the R package Gviz (Hahne and Ivanek, 2016).

#### Plasma analysis

Blood samples were collected from the tail tip of 4 h-fasted mice using chilled paraoxon-coated capillaries. sMR serum levels were determined after immune precipitation using a MR-specific antibody, followed by fluorimetry. Blood glucose levels were determined using a Glucometer (Accu-Check, Roche Diagnostics). Plasma total cholesterol (TC; Instruchemie), triglycerides (TG; Instruchemie) and insulin (Chrystal Chem) were determined using commercially available kits according to the manufacturer’s instructions. The homeostatic model assessment of insulin resistance (HOMA-IR) adapted to mice was calculated as ([glucose (mg/dl)*0.055] × [insulin (ng/ml) × 172.1])/3857 and used as a surrogate measure of whole-body insulin resistance (Lee et al., 2008). Plasma alanine aminotransferase (ALAT) was measured using a Reflotron® kit (Roche diagnostics).

#### Insulin and glucose tolerance tests

Body composition was measured by MRI using an EchoMRI (Echo Medical Systems). Wholebody insulin sensitivity was assessed after the indicated time on CD or HFD in 4 h fasted mice by an i.p. insulin tolerance test (ipITT). After an initial blood collection (t=0), an i.p. bolus of insulin (1 U/kg lean body mass; NOVORAPID, Novo Nordisk) was administered to the mice. Blood glucose was measured by tail bleeding at 15, 30, 60, and 120 min after insulin administration using a Glucometer. Whole-body glucose tolerance was assessed after the indicated time on CD or HFD in 6 h fasted mice by an intraperitoneal glucose tolerance test (ipGTT). After an initial blood collection (t=0), an i.p. injection of glucose (2g D-Glucose/kg total body weight, Sigma-Aldrich) was administered in conscious mice. Blood glucose was measured by tail bleeding at 15, 30, 60 and 120 min after glucose administration using a Glucometer (Accu-Check, Roche Diagnostics). At 15 minutes, blood was also collected for analysis of plasma insulin levels as described above.

#### Histological analyses

A piece of liver (~30 mg) was fixed in 4% paraformaldehyde (PFA; Sigma-Aldrich), paraffin-embedded, sectioned at 4 μm and stained with Hematoxylin and Eosin (H&E). Six fields at 20x magnification (total area 1.68 mm^2^) were used for the analysis of hepatic steatosis.

#### Hepatic lipid composition

Liver lipids were extracted as previously described (Geerling et al., 2014). Briefly, liver samples (~50 mg) were homogenized in ice-cold methanol. After centrifugation, lipids were extracted with CH3OH:CHCl3 (1:3 v/v), followed by phase separation with centrifugation (13,000 x g; 15 min at RT). The organic phase was dried and dissolved in 2% Triton X-100 in water. Triglycerides (TG), total cholesterol (TC) and phospholipids (PL) concentrations were measured using commercial kits (Roche Molecular Biochemicals). Liver lipids were expressed as nanomoles per mg protein, which was determined using the Bradford protein assay kit (Sigma-Aldrich).

#### RNA purification and qRT-PCR

RNA was extracted from snap-frozen tissue samples using Tripure RNA Isolation reagent (Roche Diagnostics). Total RNA (1 μg) was reverse transcribed and quantitative real-time PCR was next performed with the SYBR Green Core Kit on a MyIQ thermal cycler (Bio-Rad). mRNA expression was normalized to *Rplp0* mRNA content and expressed as fold change compared to wild-type CD-fed mice using the ΔΔCT method.

#### Isolation of stromal vascular fraction from adipose tissue

Epididymal adipose tissues were collected in PBS, then minced and digested for 1 h at 37°C in HEPES-buffered Krebs solution (pH 7.4) containing 0.5 mg/mL collagenase type I from *Clostridium histolyticum* (Sigma-Aldrich), 2% (w/v) bovine serum albumin (BSA, fraction V, Sigma-Aldrich) and 6 mM glucose. Disaggregated adipose tissues were passed through 100 μm cell strainers or 200 μm filters and washed with PBS supplemented with 2.5 mM EDTA and 5% FCS. Filtrate was either directly centrifuged (350 x *g*, 10 min at RT) or rested for 10 minutes, after which infranatant was collected and centrifuged. After centrifugation, the supernatant was discarded and the pellet containing the stromal vascular fraction (SVF) was treated with erythrocyte lysis buffer (BD Biosciences). After washing and manual counting, cells were stained with the live/dead marker Aqua or Zombie-UV (Invitrogen), fixed with 1.9% paraformaldehyde (Sigma-Aldrich) and stored in FACS buffer (PBS, 2 mM EDTA, 0.5% BSA [w/v]) at 4 °C in the dark until subsequent analysis performed within 4 days.

#### Isolation of leukocytes from liver tissue

Livers were collected in ice-cold RPMI 1640 + Glutamax (Life Technologies), minced and digested for 45 min at 37 °C in RPMI 1640 + Glutamax supplemented with 1 mg/mL collagenase type IV from *Clostridium histolyticum*, 2000 U/mL DNase (both Sigma-Aldrich) and 1 mM CaCl2. The digested tissues were next passed through 100 μm cell strainers and washed with PBS/EDTA/FCS. Following centrifugation (530 x g, 10 min at 4 °C), the cell pellet was resuspended in PBS/EDTA/FCS and centrifuged at 50 x g to pellet hepatocytes (3 minutes at 4 °C). The supernatant was next collected and pelleted (530 x g, 10 min at 4 °C), followed by treatment with erythrocyte lysis buffer. After washing, either macrophages or total leukocytes were isolated using LS columns and F4/80 or CD45 MicroBeads (35 μl beads per liver; Miltenyi Biotec), respectively, according to the manufacturer’s protocol. Isolated CD45^+^ cells were counted and processed for flow cytometry as for SVF, and F4/80^+^ cells were stimulated with FcMR and LPS as described above.

#### Isolation of peritoneal macrophages

Peritoneal wash was collected in PBS supplemented with 2 mM EDTA and centrifuged (530 x g, 10 min at 4 °C). Cell pellet was treated with erythrocyte lysis buffer, counted and processed for flow cytometry as for SVF, or macrophages were isolated using MS columns and F4/80 microbeads according to the manufacturer’s protocol.

#### Isolation of splenic macrophages

Spleens were collected in ice-cold RPMI 1640 + Glutamax, minced and digested for 20 min at 37 °C in RPMI 1640 + Glutamax supplemented with 1 mg/mL collagenase D (Sigma) and 2000 U/mL DNase. The digested tissues were next passed through 100 μm cell strainers and washed with PBS/EDTA/FCS. Following centrifugation (530 x g, 10 min at 4 °C), the cell pellet was treated with erythrocyte lysis buffer. Cells were either counted and processed for flow cytometry as for SVF, or macrophages were isolated using MS columns and F4/80 microbeads according to the manufacturer’s protocol.

#### Flow cytometry of WAT and liver myeloid subsets

Part of the cells were first permeabilized with 0.5% saponin (Sigma-Aldrich) or eBioscience permeabilization buffer (Invitrogen). After washing, cells were next stained with antibodies directed against CD45, CD45RA, CD45RB, CD45RC, CD45.2, CD64, Siglec-F, CD11b, Ly6C, F4/80 and CD11c in permeabilization buffer. Cells were measured on a FACSCanto flow cytometer or LSR-II (both BD Biosciences), and gates were set according to Fluorescence Minus One (FMO) controls.

### Quantification and statistical analysis

All *in vivo* data were analyzed by two-way ANOVA with multiple comparisons followed by post-hoc Fisher’s LSD test. For all other experiments, P values and statistical significance was calculated by student’s t-test. * P<0.05; ** P<0.01 and *** P<0.001. All graphs depict mean value ± SEM of at least three independent experiments.

### Data and Software availability

The complete RNAseq analysis incl. code and count data can be found under jsschrepping/Embgenbroich_2020 at https://github.com/schultzelab. Additionally, the unprocessed RNA-seq data is available online in the Gene Expression Omnibus (GEO) database (https://www.ncbi.nlm.nih.gov) under accession number GSE145369.

## References

Abu-Amer, Y., Ross, F.P., McHugh, K.P., Livolsi, A., Peyron, J.F., and Teitelbaum, S.L. (1998). Tumor necrosis factor-alpha activation of nuclear transcription factor-kappaB in marrow macrophages is mediated by c-Src tyrosine phosphorylation of Ikappa Balpha. The Journal of biological chemistry 273, 29417–29423. https://doi.org/10.1074/jbc.273.45.29417.

Andersen, E.S., Rødgaard-Hansen, S., Moessner, B., Christensen, P.B., Møller, H.J., and Weis, N. (2014). Macrophage-related serum biomarkers soluble CD163 (sCD163) and soluble mannose receptor (sMR) to differentiate mild liver fibrosis from cirrhosis in patients with chronic hepatitis C: a pilot study. European journal of clinical microbiology & infectious diseases : official publication of the European Society of Clinical Microbiology 33, 117–122. https://doi.org/10.1007/s10096-013-1936-3.

Bai, D., Ueno, L., and Vogt, P.K. (2009). Akt-mediated regulation of NFkappaB and the essentialness of NFkappaB for the oncogenicity of PI3K and Akt. International journal of cancer 125, 2863–2870. https://doi.org/10.1002/ijc.24748.

Bray, N.L., Pimentel, H., Melsted, P., and Pachter, L. (2016). Near-optimal probabilistic RNA-seq quantification. Nature biotechnology 34, 525–527. https://doi.org/10.1038/nbt.3519.

Brestoff, J.R., and Artis, D. (2015). Immune regulation of metabolic homeostasis in health and disease. Cell 161, 146–160. https://doi.org/10.1016/j.cell.2015.02.022.

Burgdorf, S., Kautz, A., Böhnert, V., Knolle, P.A., and Kurts, C. (2007). Distinct pathways of antigen uptake and intracellular routing in CD4 and CD8 T cell activation. Science (New York, N.Y.) 316, 612–616. https://doi.org/10.1126/science.1137971.

Burgdorf, S., Lukacs-Kornek, V., and Kurts, C. (2006). The mannose receptor mediates uptake of soluble but not of cell-associated antigen for cross-presentation. Journal of immunology (Baltimore, Md. : 1950) 176, 6770–6776. https://doi.org/10.4049/jimmunol.176.11.6770.

Chen, J. (2010). The Src/PI3K/Akt signal pathway may play a key role in decreased drug efficacy in obesity-associated cancer. Journal of cellular biochemistry 110, 279–280. https://doi.org/10.1002/jcb.22572.

Cheng, J., Phong, B., Wilson, D.C., Hirsch, R., and Kane, L.P. (2011). Akt fine-tunes NF-κB-dependent gene expression during T cell activation. The Journal of biological chemistry 286, 36076–36085. https://doi.org/10.1074/jbc.M111.259549.

Ding, D., Song, Y., Yao, Y., and Zhang, S. (2017). Preoperative serum macrophage activated biomarkers soluble mannose receptor (sMR) and soluble haemoglobin scavenger receptor (sCD163), as novel markers for the diagnosis and prognosis of gastric cancer. Oncology letters 14, 2982–2990. https://doi.org/10.3892/ol.2017.6547.

Doyle, S.L., Jefferies, C.A., and O’Neill, L.A. (2005). Bruton’s tyrosine kinase is involved in p65-mediated transactivation and phosphorylation of p65 on serine 536 during NFkappaB activation by lipopolysaccharide. The Journal of biological chemistry 280, 23496–23501. https://doi.org/10.1074/jbc.C500053200.

Geerling, J.J., Boon, M.R., van der Zon, G.C., van den Berg, S.A.A., van den Hoek, A.M., Lombès, M., Princen, H.M.G., Havekes, L.M., Rensen, P.C.N., and Guigas, B. (2014). Metformin lowers plasma triglycerides by promoting VLDL-triglyceride clearance by brown adipose tissue in mice. Diabetes 63, 880–891. https://doi.org/10.2337/db13-0194.

Hahne, F., and Ivanek, R. (2016). Visualizing Genomic Data Using Gviz and Bioconductor. Methods in molecular biology (Clifton, N.J.) 1418, 335–351. https://doi.org/10.1007/978-1-4939-3578-9_16.

Hotamisligil, G.S. (2017). Inflammation, metaflammation and immunometabolic disorders. Nature 542, 177–185. https://doi.org/10.1038/nature21363.

Hotamisligil, G.S., Murray, D.L., Choy, L.N., and Spiegelman, B.M. (1994). Tumor necrosis factor alpha inhibits signaling from the insulin receptor. Proceedings of the National Academy of Sciences of the United States of America 91, 4854–4858. https://doi.org/10.1073/pnas.91.11.4854.

Hussaarts, L., Smits, H.H., Schramm, G., van der Ham, A.J., van der Zon, G.C., Haas, H., Guigas, B., and Yazdanbakhsh, M. (2013). Rapamycin and omega-1: mTOR-dependent and - independent Th2 skewing by human dendritic cells. Immunology and cell biology 91, 486–489. https://doi.org/10.1038/icb.2013.31.

Ignatiadis, N., Klaus, B., Zaugg, J.B., and Huber, W. (2016). Data-driven hypothesis weighting increases detection power in genome-scale multiple testing. Nature methods 13, 577–580. https://doi.org/10.1038/nmeth.3885.

Jaitin, D.A., Adlung, L., Thaiss, C.A., Weiner, A., Li, B., Descamps, H., Lundgren, P., Bleriot, C., Liu, Z., and Deczkowska, A., et al. (2019). Lipid-Associated Macrophages Control Metabolic Homeostasis in a Trem2-Dependent Manner. Cell 178, 686–698.e14. https://doi.org/10.1016/j.cell.2019.05.054.

Jordens, R., Thompson, A., Amons, R., and Koning, F. (1999). Human dendritic cells shed a functional, soluble form of the mannose receptor. International immunology 11, 1775–1780. https://doi.org/10.1093/intimm/11.11.1775.

Kim, D., Paggi, J.M., Park, C., Bennett, C., and Salzberg, S.L. (2019). Graph-based genome alignment and genotyping with HISAT2 and HISAT-genotype. Nature biotechnology 37, 907–915. https://doi.org/10.1038/s41587-019-0201-4.

Klotz, L., Hucke, S., Thimm, D., Classen, S., Gaarz, A., Schultze, J., Edenhofer, F., Kurts, C., Klockgether, T., and Limmer, A., et al. (2009). Increased antigen cross-presentation but impaired cross-priming after activation of peroxisome proliferator-activated receptor gamma is mediated by up-regulation of B7H1. Journal of immunology (Baltimore, Md. : 1950) 183, 129–136. https://doi.org/10.4049/jimmunol.0804260.

Kreer, C., Kuepper, J.M., Zehner, M., Quast, T., Kolanus, W., Schumak, B., and Burgdorf, S. (2017). N-glycosylation converts non-glycoproteins into mannose receptor ligands and reveals antigen-specific T cell responses in vivo. Oncotarget 8, 6857–6872. https://doi.org/10.18632/oncotarget.14314.

Lackey, D.E., and Olefsky, J.M. (2016). Regulation of metabolism by the innate immune system. Nature reviews. Endocrinology 12, 15–28. https://doi.org/10.1038/nrendo.2015.189.

Lanthier, N., Molendi-Coste, O., Horsmans, Y., van Rooijen, N., Cani, P.D., and Leclercq, I.A. (2010). Kupffer cell activation is a causal factor for hepatic insulin resistance. American journal of physiology. Gastrointestinal and liver physiology 298, G107–16. https://doi.org/10.1152/ajpgi.00391.2009.

Lee, S., Muniyappa, R., Yan, X., Chen, H., Yue, L.Q., Hong, E.-G., Kim, J.K., and Quon, M.J. (2008). Comparison between surrogate indexes of insulin sensitivity and resistance and hyperinsulinemic euglycemic clamp estimates in mice. American journal of physiology. Endocrinology and metabolism 294, E261–70. https://doi.org/10.1152/ajpendo.00676.2007.

Lee, S.J., Evers, S., Roeder, D., Parlow, A.F., Risteli, J., Risteli, L., Lee, Y.C., Feizi, T., Langen, H., and Nussenzweig, M.C. (2002). Mannose receptor-mediated regulation of serum glycoprotein homeostasis. Science (New York, N.Y.) 295, 1898–1901. https://doi.org/10.1126/science.1069540.

Liberzon, A., Subramanian, A., Pinchback, R., Thorvaldsdóttir, H., Tamayo, P., and Mesirov, J.P. (2011). Molecular signatures database (MSigDB) 3.0. Bioinformatics (Oxford, England) 27, 1739–1740. https://doi.org/10.1093/bioinformatics/btr260.

Lluis, J.M., Buricchi, F., Chiarugi, P., Morales, A., and Fernandez-Checa, J.C. (2007). Dual role of mitochondrial reactive oxygen species in hypoxia signaling: activation of nuclear factor-{kappa}B via c-SRC and oxidant-dependent cell death. Cancer research 67, 7368–7377. https://doi.org/10.1158/0008-5472.CAN-07-0515.

Loonen, A.J.M., Leijtens, S., Serin, O., Hilbink, M., Wever, P.C., van den Brule, A.J.C., and Toonen, E.J.M. (2019). Soluble mannose receptor levels in blood correlate to disease severity in patients with community-acquired pneumonia. Immunology letters 206, 28–32. https://doi.org/10.1016/j.imlet.2018.12.001.

Love, M.I., Huber, W., and Anders, S. (2014). Moderated estimation of fold change and dispersion for RNA-seq data with DESeq2. Genome biology 15, 550. https://doi.org/10.1186/s13059-014-0550-8.

Lumeng, C.N., Bodzin, J.L., and Saltiel, A.R. (2007). Obesity induces a phenotypic switch in adipose tissue macrophage polarization. The Journal of clinical investigation 117, 175–184. https://doi.org/10.1172/JCI29881.

Martinez-Pomares, L. (2012). The mannose receptor. Journal of leukocyte biology 92, 1177–1186. https://doi.org/10.1189/jlb.0512231.

Martinez-Pomares, L., Wienke, D., Stillion, R., McKenzie, E.J., Arnold, J.N., Harris, J., McGreal, E., Sim, R.B., Isacke, C.M., and Gordon, S. (2006). Carbohydrate-independent recognition of collagens by the macrophage mannose receptor. European journal of immunology 36, 1074–1082. https://doi.org/10.1002/eji.200535685.

Martínez-Pomares, L., Crocker, P.R., Da Silva, R., Holmes, N., Colominas, C., Rudd, P., Dwek, R., and Gordon, S. (1999). Cell-specific glycoforms of sialoadhesin and CD45 are counter-receptors for the cysteine-rich domain of the mannose receptor. The Journal of biological chemistry 274, 35211–35218. https://doi.org/10.1074/jbc.274.49.35211.

Martínez-Pomares, L., Mahoney, J.A., Káposzta, R., Linehan, S.A., Stahl, P.D., and Gordon, S. (1998). A functional soluble form of the murine mannose receptor is produced by macrophages in vitro and is present in mouse serum. The Journal of biological chemistry 273, 23376–23380. https://doi.org/10.1074/jbc.273.36.23376.

Morinaga, H., Mayoral, R., Heinrichsdorff, J., Osborn, O., Franck, N., Hah, N., Walenta, E., Bandyopadhyay, G., Pessentheiner, A.R., and Chi, T.J., et al. (2015). Characterization of distinct subpopulations of hepatic macrophages in HFD/obese mice. Diabetes 64, 1120–1130. https://doi.org/10.2337/db14-1238.

Neuschwander-Tetri, B.A. (2010). Hepatic lipotoxicity and the pathogenesis of nonalcoholic steatohepatitis: the central role of nontriglyceride fatty acid metabolites. Hepatology (Baltimore, Md.) 52, 774–788. https://doi.org/10.1002/hep.23719.

Pilling, D., Fan, T., Huang, D., Kaul, B., and Gomer, R.H. (2009). Identification of markers that distinguish monocyte-derived fibrocytes from monocytes, macrophages, and fibroblasts. PloS one 4, e7475. https://doi.org/10.1371/journal.pone.0007475.

Rauen, J., Kreer, C., Paillard, A., van Duikeren, S., Benckhuijsen, W.E., Camps, M.G., Valentijn, A.R.P.M., Ossendorp, F., Drijfhout, J.W., and Arens, R., et al. (2014). Enhanced cross-presentation and improved CD8+ T cell responses after mannosylation of synthetic long peptides in mice. PloS one 9, e103755. https://doi.org/10.1371/journal.pone.0103755.

Rødgaard-Hansen, S., Rafique, A., Weis, N., Wejse, C., Nielsen, H., Pedersen, S.S., Møller, H.J., and Kronborg, G. (2015). Increased concentrations of the soluble mannose receptor in serum from patients with pneumococcal bacteraemia, and prediction of survival. Infectious diseases (London, England) 47, 203–208. https://doi.org/10.3109/00365548.2014.984321.

Saha, B., Tornai, D., Kodys, K., Adejumo, A., Lowe, P., McClain, C., Mitchell, M., McCullough, A., Dasarathy, S., and Kroll-Desrosiers, A., et al. (2019). Biomarkers of Macrophage Activation and Immune Danger Signals Predict Clinical Outcomes in Alcoholic Hepatitis. Hepatology (Baltimore, Md.) 70, 1134–1149. https://doi.org/10.1002/hep.30617.

Schuette, V., Embgenbroich, M., Ulas, T., Welz, M., Schulte-Schrepping, J., Draffehn, A.M., Quast, T., Koch, K., Nehring, M., and König, J., et al. (2016). Mannose receptor induces T-cell tolerance via inhibition of CD45 and up-regulation of CTLA-4. Proceedings of the National Academy of Sciences of the United States of America 113, 10649–10654. https://doi.org/10.1073/pnas.1605885113.

Shannon, P., Markiel, A., Ozier, O., Baliga, N.S., Wang, J.T., Ramage, D., Amin, N., Schwikowski, B., and Ideker, T. (2003). Cytoscape: a software environment for integrated models of biomolecular interaction networks. Genome research 13, 2498–2504. https://doi.org/10.1101/gr.1239303.

Shrivastava, P., Katagiri, T., Ogimoto, M., Mizuno, K., and Yakura, H. (2004). Dynamic regulation of Src-family kinases by CD45 in B cells. Blood 103, 1425–1432. https://doi.org/10.1182/blood-2003-03-0716.

Shulman, G.I. (2014). Ectopic fat in insulin resistance, dyslipidemia, and cardiometabolic disease. The New England journal of medicine 371, 2237–2238. https://doi.org/10.1056/NEJMc1412427.

Soneson, C., Love, M.I., and Robinson, M.D. (2015). Differential analyses for RNA-seq: transcript-level estimates improve gene-level inferences. F1000Research 4, 1521. https://doi.org/10.12688/f1000research.7563.2.

Suzuki, Y., Shirai, M., Asada, K., Yasui, H., Karayama, M., Hozumi, H., Furuhashi, K., Enomoto, N., Fujisawa, T., and Nakamura, Y., et al. (2018). Macrophage mannose receptor, CD206, predict prognosis in patients with pulmonary tuberculosis. Scientific reports 8, 13129. https://doi.org/10.1038/s41598-018-31565-5.

Takahashi, K., Donovan, M.J., Rogers, R.A., and Ezekowitz, R.A. (1998). Distribution of murine mannose receptor expression from early embryogenesis through to adulthood. Cell and tissue research 292, 311–323. https://doi.org/10.1007/s004410051062.

van Vliet, S.J., Gringhuis, S.I., Geijtenbeek, T.B.H., and van Kooyk, Y. (2006). Regulation of effector T cells by antigen-presenting cells via interaction of the C-type lectin MGL with CD45. Nature immunology 7, 1200–1208. https://doi.org/10.1038/ni1390.

Weber, A.N.R., Bittner, Z., Liu, X., Dang, T.-M., Radsak, M.P., and Brunner, C. (2017). Bruton’s Tyrosine Kinase: An Emerging Key Player in Innate Immunity. Frontiers in immunology 8, 1454. https://doi.org/10.3389/fimmu.2017.01454.

Westcott, D.J., Delproposto, J.B., Geletka, L.M., Wang, T., Singer, K., Saltiel, A.R., and Lumeng, C.N. (2009). MGL1 promotes adipose tissue inflammation and insulin resistance by regulating 7/4hi monocytes in obesity. The Journal of experimental medicine 206, 3143–3156. https://doi.org/10.1084/jem.20091333.

Wijngaarden, M.A., van der Zon, G.C., van Dijk, K.W., Pijl, H., and Guigas, B. (2013). Effects of prolonged fasting on AMPK signaling, gene expression, and mitochondrial respiratory chain content in skeletal muscle from lean and obese individuals. American journal of physiology. Endocrinology and metabolism 304, E1012–21. https://doi.org/10.1152/ajpendo.00008.2013.

Xie, X., Lan, T., Chang, X., Huang, K., Huang, J., Wang, S., Chen, C., Shen, X., Liu, P., and Huang, H. (2013). Connexin43 mediates NF-κB signalling activation induced by high glucose in GMCs: involvement of c-Src. Cell communication and signaling : CCS 11, 38. https://doi.org/10.1186/1478-811X-11-38.

Xue, J., Schmidt, S.V., Sander, J., Draffehn, A., Krebs, W., Quester, I., Nardo, D. de, Gohel, T.D., Emde, M., and Schmidleithner, L., et al. (2014). Transcriptome-based network analysis reveals a spectrum model of human macrophage activation. Immunity 40, 274–288. https://doi.org/10.1016/j.immuni.2014.01.006.

Yu, G., Wang, L.-G., Han, Y., and He, Q.-Y. (2012). clusterProfiler: an R package for comparing biological themes among gene clusters. Omics : a journal of integrative biology 16, 284–287. https://doi.org/10.1089/omi.2011.0118.

