## Supplemental Files for "Soluble mannose receptor induces pro-inflammatory macrophage activation and metaflammation"

***Supplementary Information***

***Supplementary Figure 1***

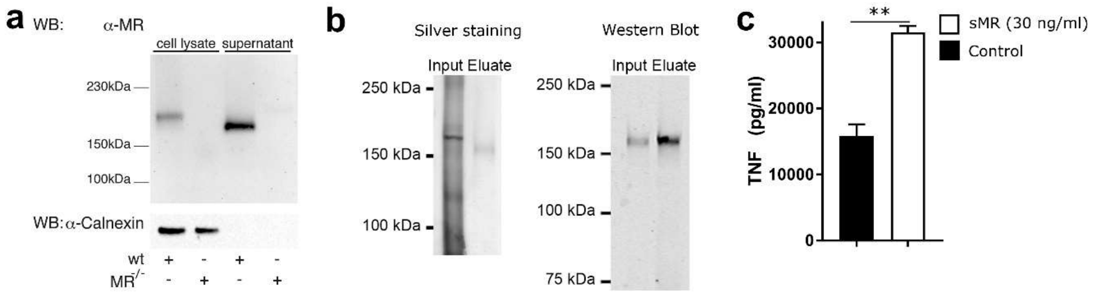


*Supplementary Figure 1: Purification of sMR from the supernatant of MR-expressing cells and its effect on cytokine secretion.*

a) The presence of the MR in the cell lysates or supernatant from wild-type or MR-deficient macrophages was depicted by Western Blot. b) sMR was purified by affinity chromatography using a Protein A/G column covalently linked to an MR-specific antibody. Images depict silver staining (left) and Western Blot using a MR-specific antibody (right). c) MR-deficient macrophages were treated with purified sMR and stimulated with LPS. Secretion of TNF was determined after 18 h by ELISA. All graphs are depicted as mean ± SEM; for all experiments, n ≥ 3.

*Supplementary Figure 2*

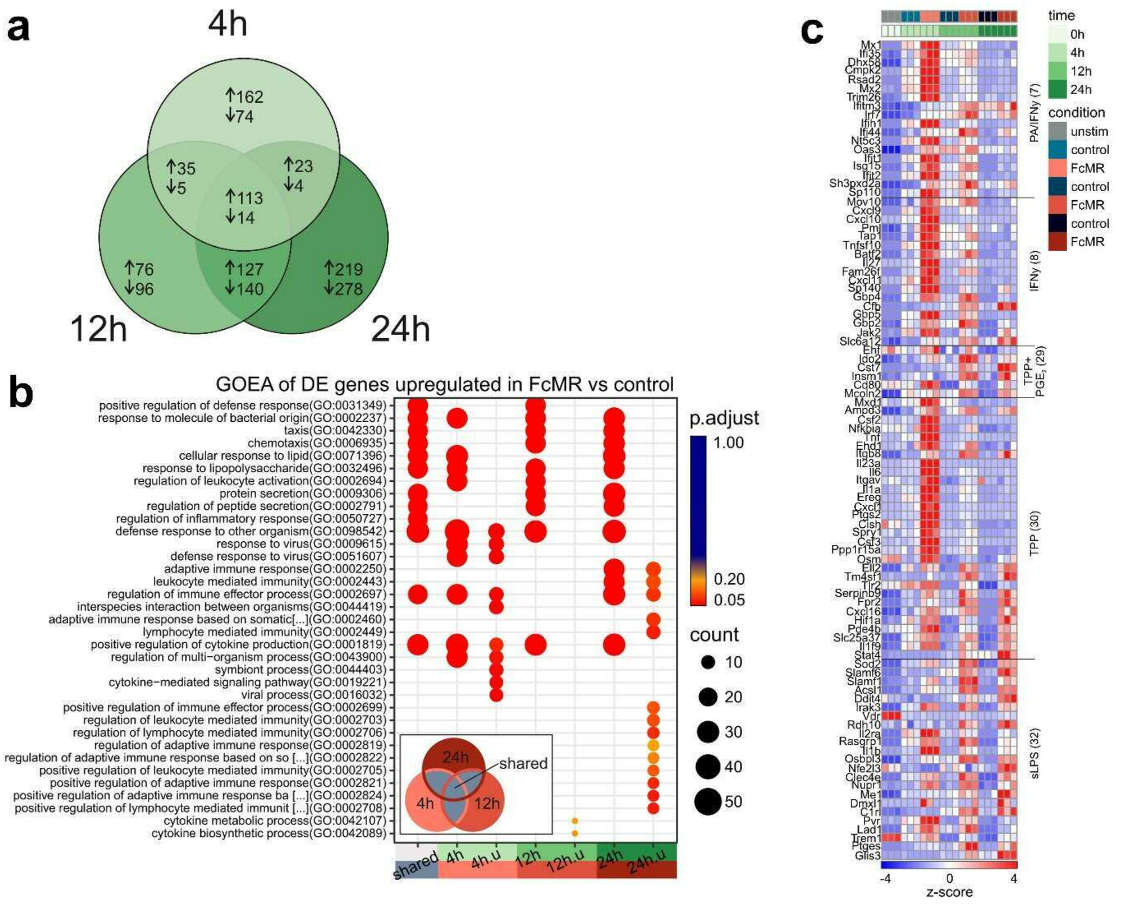


*Supplementary Figure 2: RNAseq analysis of bone marrow-derived macrophages incubated with FcMR or isotype control for 4h, 12h or 24h.*

a) Venn diagram of 1366 DE genes between FcMR-treated and control samples. b) Dot plot of gene ontology enrichment analysis (GOEA) results of the DE genes shared between all three DE gene sets (shared), the respective DE gene sets for each time point (4 h, 12 h and 24 h) and those DE genes unique for each time point (4 h.u, 12 h.u and 24 h.u). c) Heatmap of hierarchically clustered, normalized and z-scaled expression values of the genes corresponding to the enriched stimulus-specific macrophage expression signatures shown in Figure 2e.

***Supplementary Figure 3***

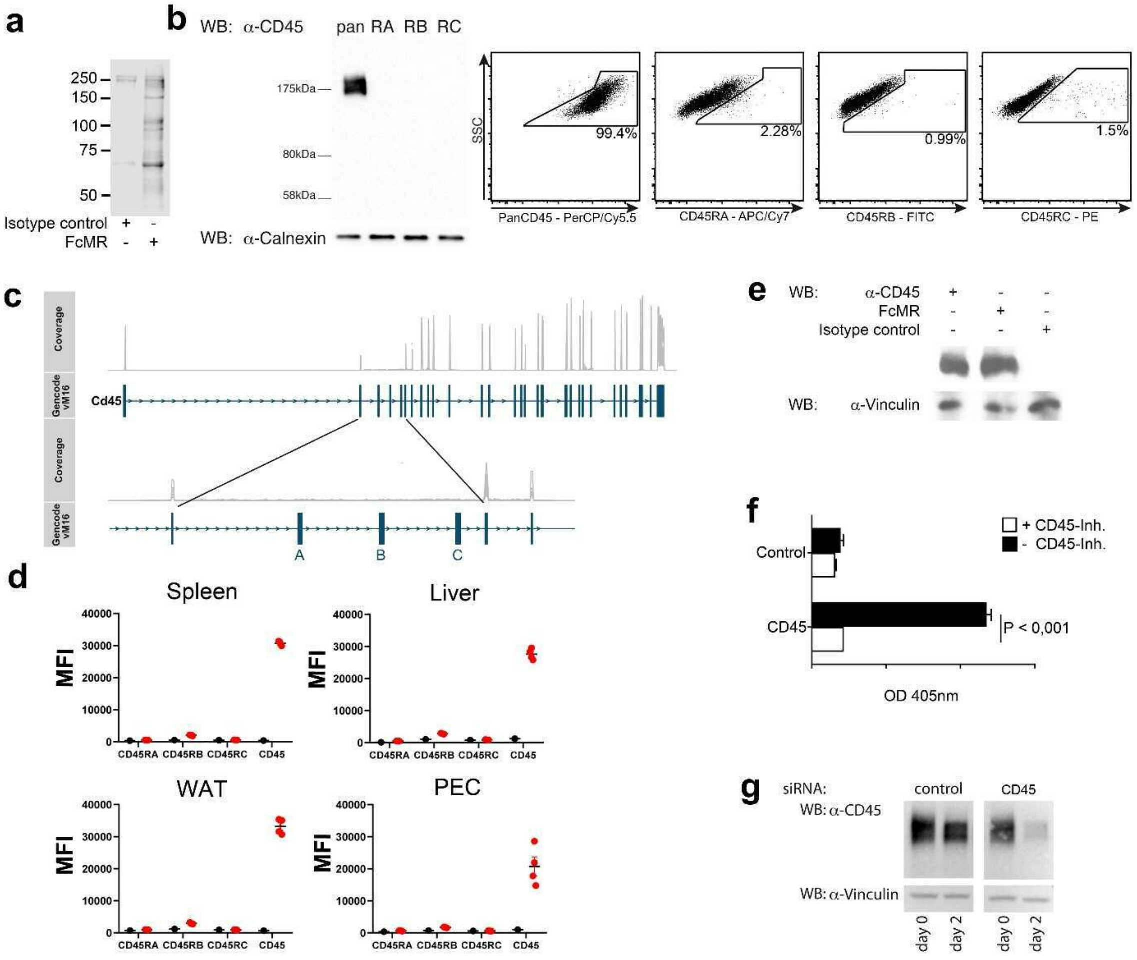


*Supplementary Figure 3: Inhibition of CD45 by the MR.*

a) Lysates from surface biotinylated macrophages were immune precipitated with FcMR or isotype control and analyzed by Western Blot using neutravidin.b) CD45 isoforms on bone marrow-derived macrophages analyzed by Western Blot and flow cytometry. c) Visualization of the RNAseq read coverage of the murine *Cd45* locus. d) Analysis of CD45 isoforms on macrophages from spleen, liver, white adipose tissue (WAT) or the peritoneal cavity (PEC) by flow cytometry. Black dots indicate FMO (fluorescence minus one) controls. e) F4/80$^{+}$ splenic macrophages were isolated by magnetic separation. Cell lysates were analyzed by far Western Blot after staining with FcMR or isotype control or by Western Blot with antibodies against

CD45 and vinculin. f) CD45 was immune precipitated from macrophage cell lysates and incubated with 4-NPP in the presence or absence of the CD45 inhibitor SF1670. Graph depicts dephosphorylation of 4-NPP. Samples without CD45 antibody were used as controls. g) siRNA-mediated down-regulation of CD45. All graphs are depicted as mean ± SEM; for all experiments, $n \geq 3$.

***Supplementary Figure 4***

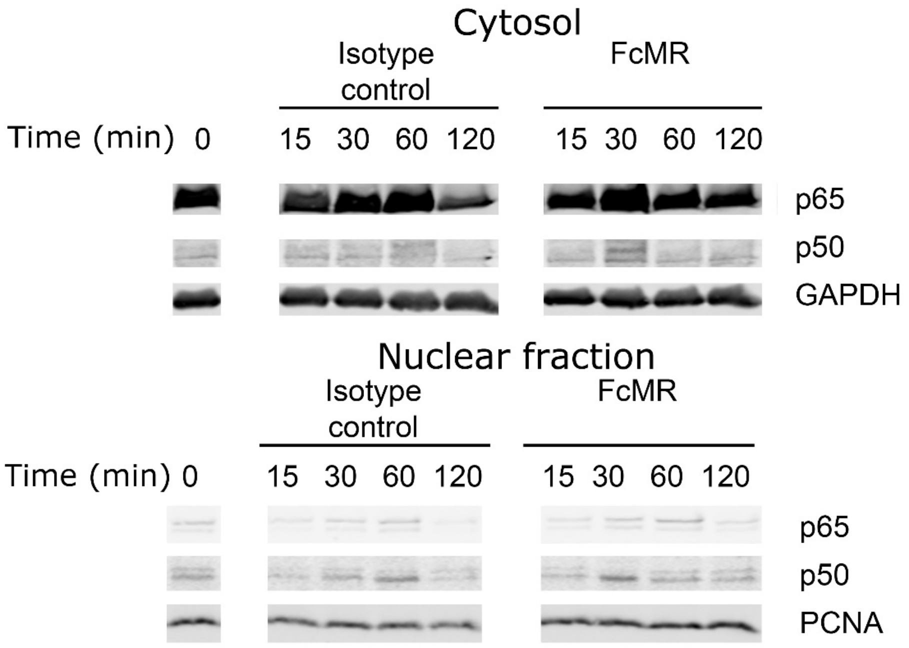


*Supplementary Figure 4: Akt inhibition blocks MR-induced nuclear translocation of p65 and p50.*

MR-deficient macrophages were incubated with FcMR or isotype control in the presence of 5 µM of the Akt inhibitor MK-2206. p65 and p50 were determined in the cytosolic and nuclear fraction by Western Blot.

*Supplementary Figure 5*

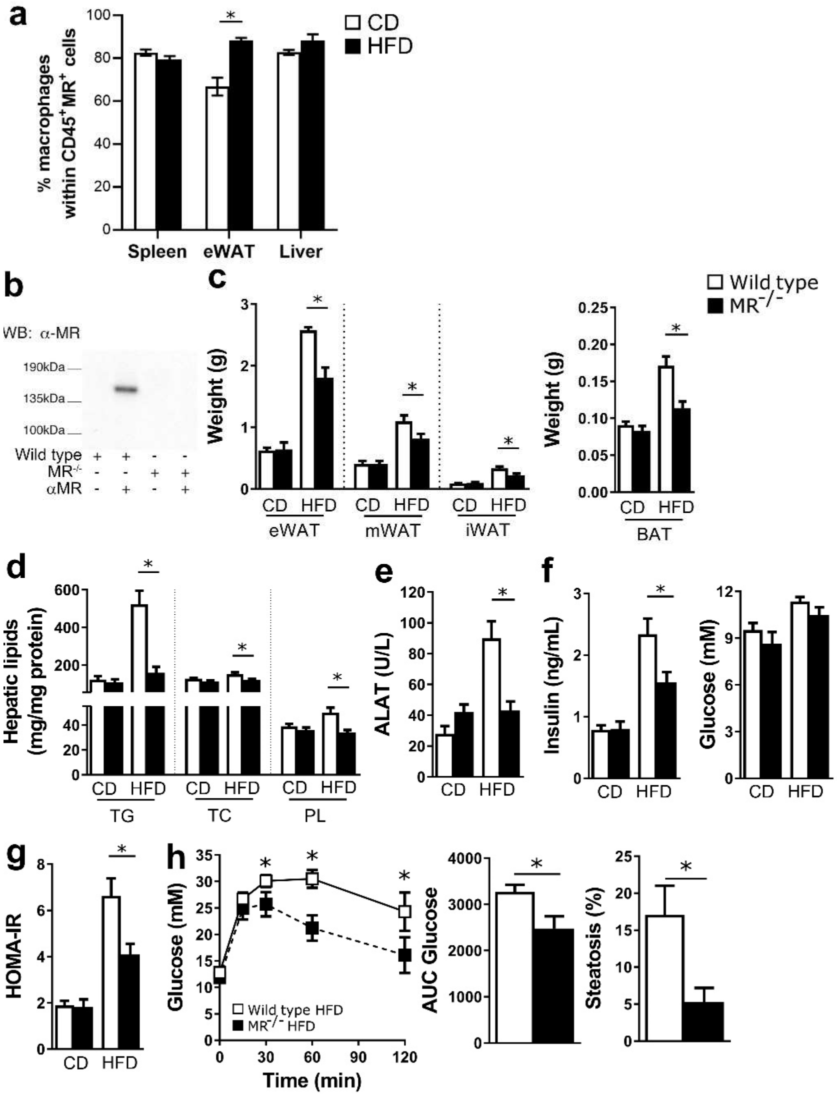


*Supplementary Figure 5: MR serum levels in obesity and effect of the MR on weight gain and whole body glucose and insulin tolerance.*

a) Percentage of F4/80$^+$/CD64$^+$ macrophages within CD45$^+$MR$^+$ cells in the indicated tissues b) sMR levels immune precipitated from the serum of wild-type or MR-deficient mice and depicted by Western Blot. c) Weight of different white fat pads or brown fat after 18 weeks of diet. d) Liver triglycerides (TG), total cholesterol (TC) and phospholipids (PL) after 18 weeks of diet. e) The plasma concentrations of alanine aminotransferase (ALAT) in pooled samples of 2-3 mice from two separate experiments (3-4 pooled samples per group). f) Plasma insulin and glucose levels after 18 weeks of diet. g) HOMA-IR in 4 h-unfed mice after 18 weeks of diet. h) Intraperitoneal glucose tolerance test and steatosis in wild-type and MR-deficient mice that were matched on body weight after 18 weeks on HFD. Results are expressed as means ± SEM; n=10-15 mice per group, except for  a (n=3 per group) and b (n=6-8 mice per group).

*Supplementary Figure 6*

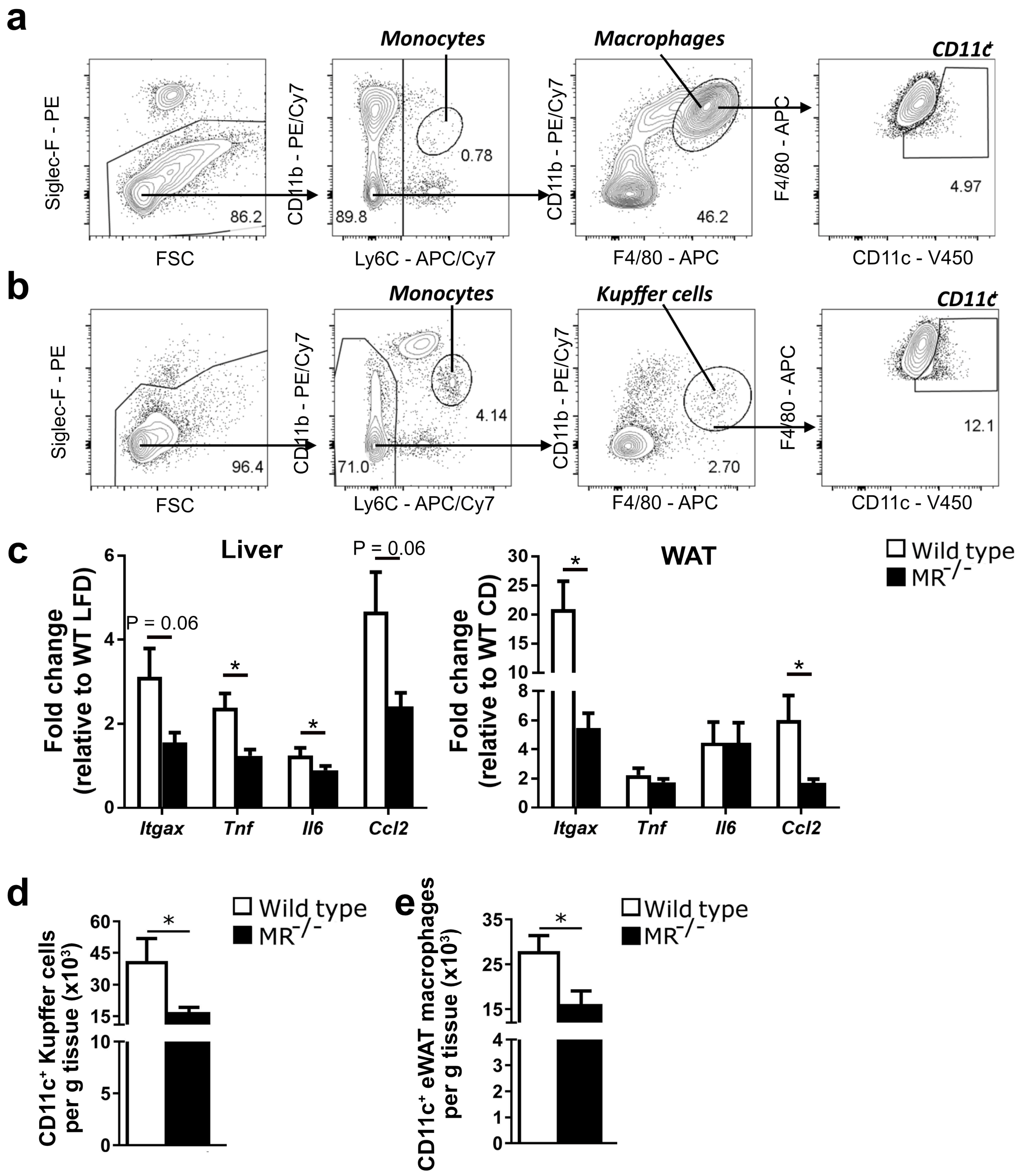


*Supplementary Figure 6: Effect of MR deficiency on inflammatory macrophages in obesity.*

a-b) Gating strategy to quantify myeloid cell populations in adipose tissue (a) and liver (b). c) mRNA expression levels of selected activation markers in liver and gonadal WAT after 18 weeks on HFD monitored by RT-qPCR (Itgax: CD11c; Tnf: TNF; Il6: IL-6; Ccl2: MCP-1).

Results are expressed as relative to the housekeeping gene Rplp0 (RPLP0/36B4) as fold change vs wild-type CD mice. d-e) Mice were matched on body weight after 18 weeks on HFD. Graphs indicate numbers of $CD11c^{+}$ macrophages in liver (d) or eWAT (e). Results are expressed as means ± SEM; n=5 mice per group.

***Supplementary Figure 7***

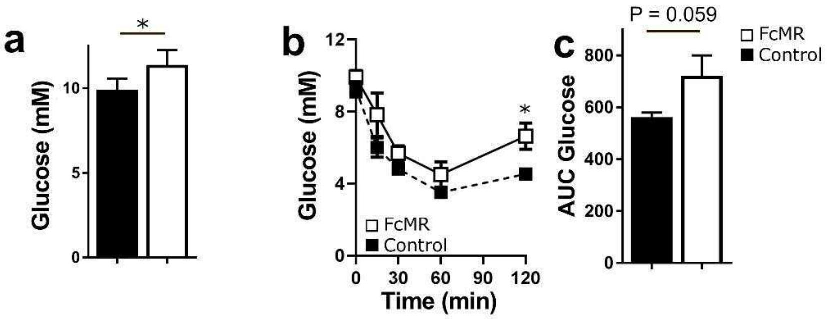


*Supplementary Figure 7: sMR treatment regulates whole body metabolism in HFD-fed mice.*

Wild-type mice were fed a HFD and concomitantly injected i.p. with 4,82 μmoles/mouse FcMR or isotype control every 3 days. After 4 weeks, fasting glucose levels were measured (a) and an intraperitoneal insulin tolerance test (b, c) was performed. Results are expressed as means ± SEM; n=6 mice per group.
